# Non-canonical inhibition in motion vision

**DOI:** 10.64898/2026.11.19.767796

**Authors:** Birte Zuidinga, Jesús Pujol-Martí, Renée M. Vieira, Eleni Samara, Cemre Coşkun, Melisa Özmen, Alexander Borst, Lukas N. Groschner

**Affiliations:** Max Planck Institute for Biological Intelligence, Circuits – Computation – Models, Martinsried, Germany; Medical University of Graz, Gottfried Schatz Research Center, Molecular Biology and Biochemistry, Graz, Austria; Department of Zoology, University of British Columbia, Vancouver, Canada; Department of Neuroscience, NYU School of Medicine, New York, NY, US

## Abstract

Artificial neural networks compute through simple weighted connections. Biological neurons, in contrast, perform complex non-linear operations that depend on the intrinsic biophysics of their receptors, which are often obscure. Here, we identify the receptor components of a divisive synapse implementing an AND-NOT computation in the motion-vision circuit of *Drosophila melanogaster*. We show that inhibitory synapses on direction-selective T4 and T5 neurons recruit both the histamine receptor Ort and the GABA_A r_eceptor Rdl, with Rdl acting as the primary inhibitory determinant of direction selectivity and visual motion-related behavior. Remarkably, Rdl is gated not only by ligand binding but also by membrane voltage. Using biophysical modeling to dissect these dual gating mechanisms, we reveal that voltage and ligand sensitivity independently sharpen the neuron’s directional tuning. These results link the unusual biophysical properties of a GABA_A r_eceptor to a fundamental neural computation and expand the functional repertoire of ionotropic receptors across nervous systems.

## Introduction

Shunting inhibition is a powerful mechanism to suppress neural activity. By vetoing excitatory input it implements a division-like computation in a single neuron,^1^ analogous to a logic AND-NOT gate.^2–5^ Motion vision is a conserved single-neuron computation that has been proposed to rely on shunting inhibition in rabbits,^2^ cats,^3^ mice,^6^ locusts,^7^ and flies.^8–11^ In *Drosophila melanogaster*, elementary motion detector neurons T4 and T5 receive luminance signals originating from neighboring points in visual space in the form of distinct neurotransmitters released by different presynaptic neurons (Figures 1A and 1B).^12–17^ When the signals arrive in a sequence consistent with visual motion in the preferred direction of the neuron, its activity is enhanced.^8,18–20^ Conversely, when signals arrive in the opposite sequence, consistent with visual motion in the null direction, activity is suppressed.^8–10,18,19^ This null-direction suppression is thought to result from GABAergic transmission onto the spatially oriented dendrites of T4 and T5 neurons vetoing the excitatory effect of acetylcholine. The receptor architecture that underlies this type of shunting inhibition is unknown.

**Figure 1.**
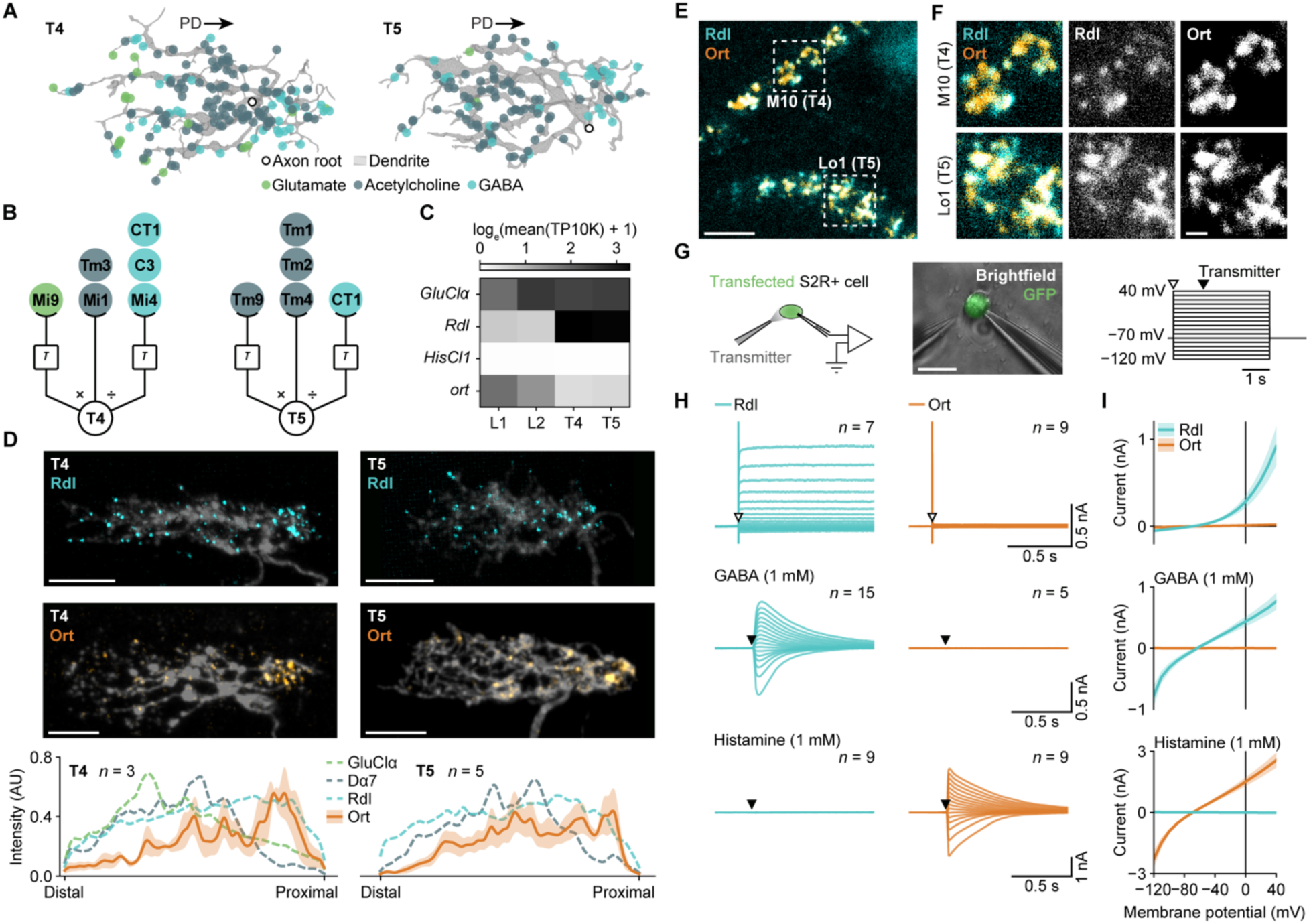
GABA and histamine receptors cluster at the proximal dendrite of direction-selective neurons. (A) T4 and T5 dendrite morphology and predicted glutamatergic (green), cholinergic (gray), and GABAergic (cyan) synapses from the FAFB v783 connectome.^14–17^ PD, preferred direction. (B) Schematic overview of the circuit and the signals resulting in direction-selective T4 and T5 neurons. The central signal is multiplied (×) and divided (÷) by temporally delayed (τ) signals from flanking ommatidia. (C) Heatmap of log1p-transformed RNA transcripts per 10,000 (TP10K) of cys-loop ligand-gated chloride channels in L1, L2, T4, and T5 neurons from single-cell RNA sequencing data (GEO: GSE156455),^27^ averaged over 24–96 hours after pupal formation. (D) Dendritic distribution of tagged receptors in representative single T4 and T5 neurons (top, Rdl;^22^ center, Ort) and average intensity profiles of all neurons (bottom). Ort data (orange) are mean ± SEM. Means of GluClα (green), Dα7 (gray), and Rdl (cyan) from Fendl et al.^22^ are shown for comparison (dashed lines). AU, arbitrary units; scale bars, 5 µm. (E) Micrograph of T4 and T5 dendrites expressing *ort::HA* (orange) and *Rdl::GFP* (cyan), representative of four optic lobes. Dashed boxes are magnified in (F) and indicate T4 dendrites in medulla layer 10 (M10) and T5 dendrites in lobula layer 1 (Lo1). Scale bar, 5 µm. (F) Magnification of marked areas in (E) (left) and separated grayscale Rdl (middle) and Ort (right) channels for M10 (top) and Lo1 (bottom). Scale bar, 1 µm. (G) Whole-cell patch-clamp recording of transfected S2R+ cells with neurotransmitter applications (left), exemplary brightfield image (middle) with native GFP signal (green), and stimulus protocol (right). Empty triangle, voltage step; filled triangle, transmitter application; scale bar, 20 µm. (H) Average currents elicited by voltage steps (top, empty triangles), applications of 1 mM GABA (center, filled triangles), and applications of 1 mM histamine (bottom, filled triangles) in cells expressing *Rdl* (cyan) or *ort* (orange) at 17 clamp potentials ranging from −120 to +40 mV. (I) Current-voltage relationships corresponding to data in (H). Data are mean ± SEM. *n* values indicate the number of cells. See also Figures S1–S4, Table S3, and STAR Methods.

Prime candidates are members of the cys-loop ligand-gated chloride channel superfamily, because the chloride reversal potential is sufficiently close to the resting potential of T4 and T5 neurons and chloride conductances are generally large enough to dissipate concurrent excitation.^20,21^ To suppress neuronal responses to motion in the null direction, possible receptors need to localize to the proximal part of the dendrite, which is the first to receive luminance signals during stimulation with null-direction motion (Figure 1A). Among the ligand-gated chloride channels of *Drosophila*, GluClα, Lcch3, and CG8916 do not localize to the proximal part of the dendrite,^22,23^ and HisCl1 and Grd are not expressed in both T4 and T5 neurons (Figures 1C and S1A).^24–27^ This leaves two ligand-gated chloride channels, encoded by the genes *Rdl* and *ort*, as possible receptors to mediate null-direction suppression. Both genes are transcribed in T4 and T5 neurons (Figures 1C and S1A).^24–27^ Even though the transcription of *ort* peaks during development, Ort protein remains detectable in T4 and T5 dendrites of adult flies.^28^ Using genetic manipulations, immunohistochemistry, electrophysiology, and two-photon imaging, we test the functional roles of Rdl and Ort receptors in motion vision.

## Results

### Subcellular distribution of GABA and histamine receptors

In single T4 and T5 neurons, *Rdl* localizes predominantly to the proximal dendrite.^22,23^ By expressing genetically tagged *ort::HA*,^29^ we found that Ort protein matched the subcellular distribution of Rdl (Figure 1D). Tagging of Rdl and Ort in the same neurons revealed clustering and partial co-localization of the two signals in dendritic regions of T4 and T5 neurons (Figures 1E and 1F). The spatial proximity of the two receptors and their asymmetric dendritic distributions suggest that Rdl and Ort play a joint role in null-direction suppression. However, while Rdl is a known GABA receptor,^30^ Ort is only described as sensitive to histamine;^31,32^ and none of the input neurons to T4 and T5 express any of the molecular machinery required for histaminergic transmission (Figure S1B).^25–27^ This prompted us to characterize the ligand specificity and the electrophysiological properties of Rdl and Ort receptors.

### Intrinsic voltage-sensitivity of a GABA_A r_eceptor

We transfected *Drosophila* S2R+ cells^33^ with control or receptor-encoding plasmids^34^ (Figure S2) and performed whole-cell voltage-clamp recordings during temporally and spatially confined neurotransmitter applications (Figure 1G). Curiously, besides their expected response to applications of 1 mM GABA, *Rdl*-expressing cells also responded to depolarizing voltage steps alone with an outwardly rectifying current that was not detectable in cells transfected with *ort* or control plasmid (Figures 1H, 1I, and S3). The *ort*-expressing cells responded as expected to applications of 1 mM histamine (Figures 1H and 1I). Transmitter-elicited currents reversed at −64.7 ± 2.8 mV for *Rdl*- and −68.7 ± 1.6 mV for *ort*-expressing cells (mean ± SEM), and were driven predominantly by chloride: Lowering the ion’s extracellular concentration from 117 mM to 14 mM shifted the average reversal potentials by 43.7 mV (*Rdl*) and 48.8 mV (*ort*) (Figure S4) which is in approximate agreement with the 53.6-mV shift expected of a pure chloride conductance.

### Lack of histamine release onto T4 and T5 neurons

The response of Ort receptors to histamine replicated previous findings,^31,32^ suggesting that Ort might confer histamine sensitivity upon T4 and T5 neurons. However, several lines of evidence from RNA sequencing, immunohistochemistry, and connectomics bear no signs of direct histaminergic input to T4 and T5 neurons: First, the presynaptic partners of T4 and T5 neurons^12,13^ express negligible amounts of gene transcripts involved in histamine synthesis (*Hdc* and *t*),^35,36^ its transmembrane transport (*lovit*),^37^ and recycling (*CarT*)^38^ compared to histaminergic photoreceptors (Figure S1B).^25–27^ Second, in the optic lobe, histamine and its vesicular transporter localize to the axon terminals of photoreceptors in the lamina and medulla but not to medulla layer 10 (M10) or lobula layer 1 (Lo1), where T4 and T5 dendrites reside, respectively.^37,39^ And third, ultrastructure-based neurotransmitter classifications for presynaptic partners of T4 and T5 neurons have minimal prediction scores for histamine.^14–17,40^

To formally rule out the possibility of histamine release onto T4 and T5 dendrites, we performed two-photon imaging of the fluorescent histamine sensor HA1m^41^ expressed simultaneously in T4 and T5 neurons as well as L1 and L2 neurons while visually stimulating the fly (Figures 2A and 2B). L1 and L2 neurons, which receive histamine from photoreceptors,^31,32,42^ served as internal positive control. In the lamina, the postsynaptic region of these neurons, HA1m signal increased in response to luminance increments and decreased in response to luminance decrements (Figure 2C), in line with the expected photoreceptor activity. This testifies that HA1m has the appropriate sensitivity to detect histamine release in the visual system. In T4 and T5 neurons of the same flies, however, the HA1m signal remained flat in response to visual stimulation with contrast edges moving in eight different directions (Figure 2C), a stimulus that elicits voltage and calcium transients in the neurons and their inputs.^11,20,43^ The absence of HA1m responses provides another case in point that native Ort on the dendrites of T4 and T5 neurons does not encounter its natural ligand histamine.

**Figure 2.**
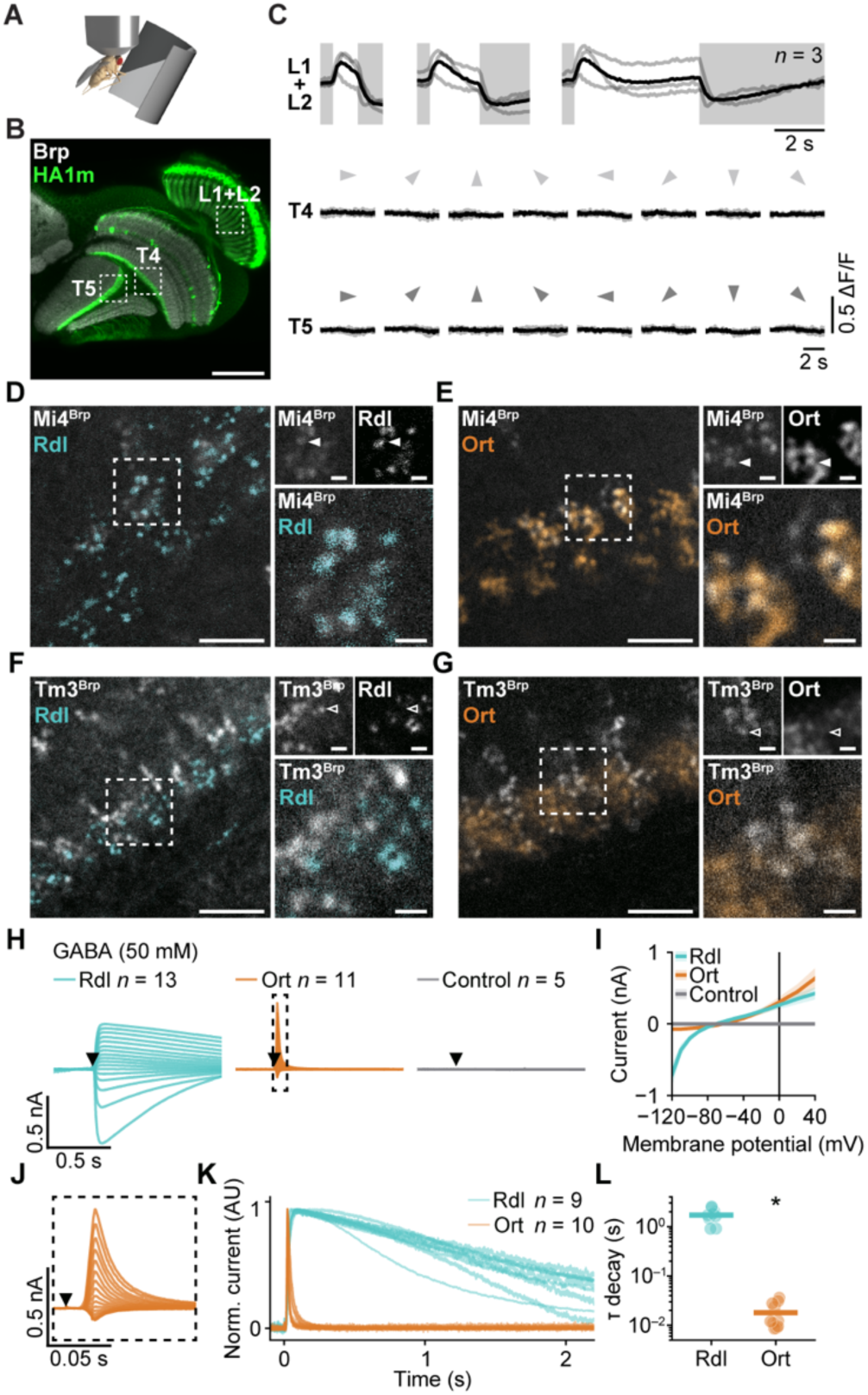
In absence of histamine release onto T4 and T5 dendrites, Rdl and Ort co-localize with GABAergic presynapses and respond to GABA. (A) Two-photon histamine imaging during visual stimulation. (B) Micrograph of an optic lobe of a fly expressing *HA1m* (green) in L1, L2, T4, and T5 neurons, counterstained with antibody against Bruchpilot (Brp, grayscale). Micrograph representative of four optic lobes. Dashed boxes indicate approximate two-photon imaging locations targeting the dendrites of L1 and L2, T4, and T5 neurons, respectively. Scale bar, 50 µm. (C) HA1m fluorescence signals of L1 and L2 dendrites in response to ON–OFF flashes (top, shaded areas represent OFF periods), T4 dendrites in response to ON edges (center), and T5 dendrites in response to OFF edges (bottom) moving at 50° s^−1^ in 8 directions (arrowheads). The light lines represent individual flies; the solid lines represent the mean. (D–G) Left, micrograph of M10 with the presynaptic marker *Brp^short^::mCherry* (grayscale) expressed in GABAergic Mi4 (D and E; Mi4^Brp^) or cholinergic Tm3 neurons (F and G; Tm3^Brp^) with *Rdl::GFP* (D and F; Rdl, cyan) or *ort::HA* (E and G; Ort, orange) expressed in T4 neurons. Right, corresponding separate channels (top, grayscale) and magnification of the dashed box on the left (bottom). Arrowheads indicate examples of co-localizing (filled) and non-co-localizing (empty) puncta. Micrographs are representative of 5 (D), 6 (E), 3 (F), and 2 (G) optic lobes. Scale bars; 5 µm (left) and 1 µm (right). (H) Average currents elicited by applications of 50 mM GABA (filled triangles) in cells expressing *Rdl* (cyan), *ort* (orange), or control plasmid (gray), at 17 clamp potentials ranging from −120 to +40 mV. The data with dashed outline are magnified in (J). (I) Current-voltage relationships corresponding to data in (H). Data are mean ± SEM. (J) Magnification of the dashed outline in (H). (K) Normalized response currents to 50 mM GABA applications at +40 mV clamp potential. The lines represent individual cells that were used for exponential decay fits (see STAR Methods). AU, arbitrary units. (L) Exponential decay time constants (τ) of individual cells in (K) (circles) and averages (horizontal lines). A significant difference in τ compared to Rdl, determined using a two-tailed Mann–Whitney *U*-test, is indicated; \**P* = 2.2 × 10^−5^ (see Table S1). *n* values indicate the number of flies (C) or cells (H and K). See also Figure S5, Tables S1 and S3, and STAR Methods.

### GABA sensitivity of a histamine receptor

The co-localization of Rdl and Ort in T4 and T5 dendrites (Figures 1E and 1F) and their structural similarity^44^ prompted us to revisit the hypothesis that both proteins bind GABA, perhaps with different sensitivity. To identify the type of synapse that the receptors associate with, we labelled presynaptic terminals of confirmed GABAergic (Mi4) and cholinergic (Tm3) neurons using a *Brp^short^::mCherry* construct^45^ and combined this with expression of labelled *Rdl::GFP*^22^ or *ort::HA*^29^ receptors in T4 neurons. Fluorescence patterns of Rdl and Ort both overlapped with GABAergic Mi4 (Figures 2D and 2E) but not with cholinergic Tm3 presynapses (Figures 2F and 2G), suggesting that they are, in fact, both in pole position for GABA exposure. Reassessment of GABA responses in *ort*-expressing S2R+ cells using a higher concentration (50 mM) revealed consistent currents in response to GABA applications (> 0.2 nA at +40 mV in 10/11 cells, Figures 2H–2J). These currents were different from those of *Rdl*-expressing cells in that they were outwardly rectified (Figures 2H–2J) and short-lived, with a decay time constant (τ) of 0.018 ± 0.010 s (mean ± SD) for *ort*-expressing cells versus 1.7 ± 0.57 s for *Rdl*-expressing cells (Figures 2K and 2l). Cells transfected with control plasmid did not exhibit GABA responses (Figures 2H and 2I), confirming that S2R+ cells have no native GABA sensitivity. The current in *ort*-expressing cells was also not caused by an unspecific effect of a high neurotransmitter concentration, since applications of 50 mM glutamate did not elicit currents in any of the conditions (Figure S5A).

To investigate the hypothesis that Rdl and Ort generate functional variability by forming heteromeric receptors, which are not uncommon among cys-loop ligand gated chloride channels,^46–49^ we transfected S2R+ cells with a plasmid encoding both genes separated by a T2A-linker. Response characteristics of cells expressing both *Rdl* and *ort* resembled those of cells expressing *Rdl* alone: GABA applications and depolarizing voltage steps gave rise to large whole-cell currents, which were absent upon stimulation with histamine (Figures S5B–S5E). A tentative explanation for the lack of a histamine response is that Ort and Rdl form heteromeric receptors lacking histamine sensitivity. This, however, might be an idiosyncrasy of the reduced system of cultured cells. The subunit composition of the receptors in T4 and T5 neurons and its influence on the neurons’ function remain unclear. To approach this question *in vivo*, we selectively knocked down each receptor from T4 and T5 neurons using RNA interference (RNAi) and characterized their visual responses using two-photon calcium imaging.

### Rdl tunes the response selectivity of T4 and T5 neurons

Immunohistochemical experiments confirmed the efficacy of gene silencing using *Rdl* RNAi^50^ and *ort* RNAi^51^ constructs. T4/T5-targeted expression of each construct significantly reduced the respective immunofluorescence in dendritic regions of T4 and T5 neurons when compared to RNAi targeting *HisCl1*^52^ (Figures 3A–3C and Table S1), another member of the cys-loop receptor family that is not expressed in T4 and T5 neurons (Figures 1C and S1A).^24–27^ For functional measurements, GCaMP8m^53^ and RNAi constructs were targeted to the subtypes a and d of T4 and T5 neurons (Figure 3D), which are sensitive to front-to-back and downward motion, respectively.^54^ To evaluate neuronal responses to stationary luminance changes, we stimulated flies with full-screen luminance increments (ON) and decrements (OFF) and measured calcium signals in dendrites of T4 (M10) and T5 neurons (Lo1), respectively. Control neurons expressing no RNAi or *HisCl1* RNAi, and those expressing *ort* RNAi, responded only weakly to these non-motion stimuli, whereas knockdown of *Rdl* alone or together with *ort* resulted in up to 3.5-fold higher response amplitudes (Figure 3E). In other words: Silencing of *Rdl* turned motion-sensitive neurons responsive to static changes in luminance.

**Figure 3.**
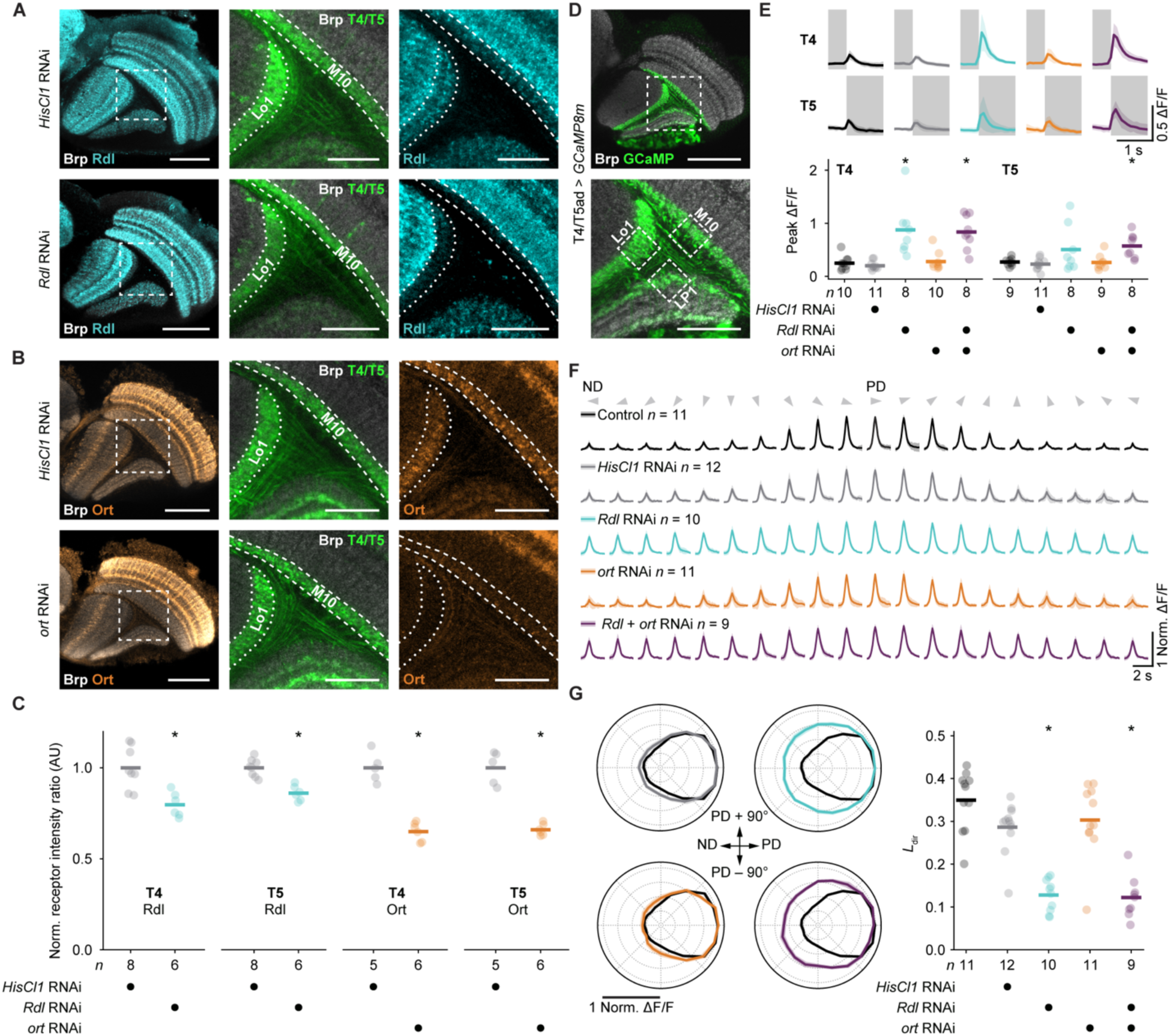
Rdl-deficient T4 and T5 neurons erroneously respond to motionless flashes and contrast edges moving in the null direction. (A) Representative Rdl immunofluorescence (cyan, left and right) in the optic lobe of flies expressing RNAi targeting *HisCl1* (top) or *Rdl* (bottom) in T4 and T5 neurons. Synaptic structures are shown in grayscale (Brp, left and middle). Dashed boxes (left) are magnified in the middle and right panels. Dendritic regions of T4 (M10, dashed lines) and T5 (Lo1, dotted lines) were identified based on anti-GFP antibody (middle, green). Scale bars, 50 µm (left) and 20 µm (middle, right). (B) Representative Ort immunofluorescence visualized as in (A), but using *ort* RNAi and anti-Ort antibody (orange). (C) Receptor fluorescence intensity ratios (see STAR Methods) in M10 (T4) and Lo1 (T5). Significant differences from *HisCl1* RNAi, determined using two-tailed Student’s *t*-tests, are indicated; \**P* = 2.5 × 10^−3^ (T4 Rdl), \**P* = 1.1 × 10^−4^ (T5 Rdl), \**P* = 1.1 × 10^−5^ (T4 Ort), and \**P* = 1.2 × 10^−5^ (T5 Ort) (see Table S1). (D) *GCaMP* expression pattern (green) driven by T4/T5ad-*GAL4*, counterstained for synaptic structures (Brp, grayscale). The dashed box (top) is magnified (bottom) to show the approximate fields of view for imaging T4ad dendrites (M10), T5ad dendrites (Lo1) and T4/T5a axon terminals (LP1). Micrograph representative of seven optic lobes. Scale bars, 50 µm (top) and 20 µm (bottom). (E) Calcium responses (mean ± SD) of T4ad (top) and T5ad neurons (center) without (control, black) or with RNAi targeting *HisCl1* (gray), *Rdl* (cyan), *ort* (orange), or *Rdl* + *ort* (purple) to ON- and OFF-flashes, respectively, and corresponding response maxima (bottom). Shaded areas indicate OFF periods. Significant response differences compared to control, determined using Dunn’s multiple comparisons test following Kruskal–Wallis test, are indicated; \**P* = 4.4 × 10^−3^ (T4, *Rdl* RNAi), \**P* = 2.2 × 10^−3^ (T4, *Rdl* + *ort* RNAi), \**P* = 4.6 × 10^−2^ (T5, *Rdl* + *ort* RNAi). (F) Normalized T4a calcium responses (mean ± SD) to ON-edges moving at 50° s^−1^ in 20 different directions (arrowheads). (G) Data from (F) represented as polar plots (mean ± SEM) of the maximum responses (left) and corresponding *L*dir values (right). No-RNAi control data (black) are repeated in each polar plot for comparison. Significant differences in *L*dir compared to no-RNAi controls, determined using Dunn’s multiple comparisons test following Kruskal–Wallis test, are indicated; \**P* = 3.5 × 10^−5^ (*Rdl* RNAi), \**P* = 2.3 × 10^−5^ (*Rdl* + *ort* RNAi). Circles represent individual flies; horizontal lines represent genotype averages. PD, preferred direction; ND, null direction*. n* values indicate the number of flies. See also Figures S6–S8, Tables S1 and S3, and STAR Methods.

To test the direction selectivity of the neurons, contrast edges were moved across the screen at 50° s^−1^ in 20 different directions separated by 18°. To capture direction-selective responses of a single subtype, we measured calcium signals in the lobula plate, where T4 and T5 subtypes segregate into separate layers. Since the characteristic ON-(T4) and OFF-selectivity (T5) was maintained in all genetic conditions (Figure S6), we could differentiate T4a and T5a neurons that both innervate lobula plate layer 1 by stimulating with single ON and OFF edges, respectively. Control neurons showed sharp directional tuning with strong response attenuation as the stimulus deviated from their preferred direction. Interference with *Rdl* expression increased responses to motion in non-preferred directions (Figure 3F), broadened the directional tuning curve, and caused a significant reduction in the directional tuning index (*L*_dir)_ compared to controls (*P* = 3.5 × 10^−5^, Kruskal–Wallis test followed by Dunn’s multiple comparisons test; Figure 3G and Table S1). Effects were similar for T4 and T5 neurons and were only observed in neurons expressing *Rdl* RNAi alone or alongside *ort* RNAi, not in neurons expressing *ort* RNAi alone (Figures 3F, 3G, and S7), speaking against a crucial role of Ort in the direction selectivity of T4 and T5 neurons. To investigate a possible function of Ort in neuronal development, we quantified the dendritic morphologies of single neurons. No significant differences in dendritic size, density, and orientation were detected upon comparing control T4 neurons with neurons expressing *ort* RNAi for all subtypes (Figure S8 and Table S2). Therefore, the function of Ort in T4 and T5 neurons—if there is any—remains unclear and Rdl emerges as the main receptor suppressing responses to global luminance changes and motion in non-preferred directions.

### Model-based separation of the voltage- and ligand-gated function of Rdl

The relative contribution of the voltage- and ligand-gated components of Rdl currents (Figure 1H) remains unclear, because they are not separable by RNAi. To isolate them nonetheless, we used a biophysically realistic, compartmentalized model of a T4 neuron. The three-arm model took into account the precise synaptic organization (Figure 1A)^12–17^ and the temporal dynamics^20,55^ of delayed glutamatergic disinhibition, fast cholinergic excitation, and slow GABAergic inhibition (see STAR Methods). The axon contained voltage-gated sodium channels^56^ and GABAergic synapses were endowed with an additional voltage-gated chloride conductance mimicking the current-voltage relationship and the gating kinetics of Rdl (Figures 1H and 1I). The model produced sharply tuned, direction-selective membrane potential responses, which broadened upon blocking of the voltage-gated component of Rdl, and lost most of their selectivity upon complete removal of Rdl (Figures 4A and 4B). This suggests that Rdl exerts its tuning effect not only through canonical ligand-gating, but also through non-canonical voltage-gating.

**Figure 4.**
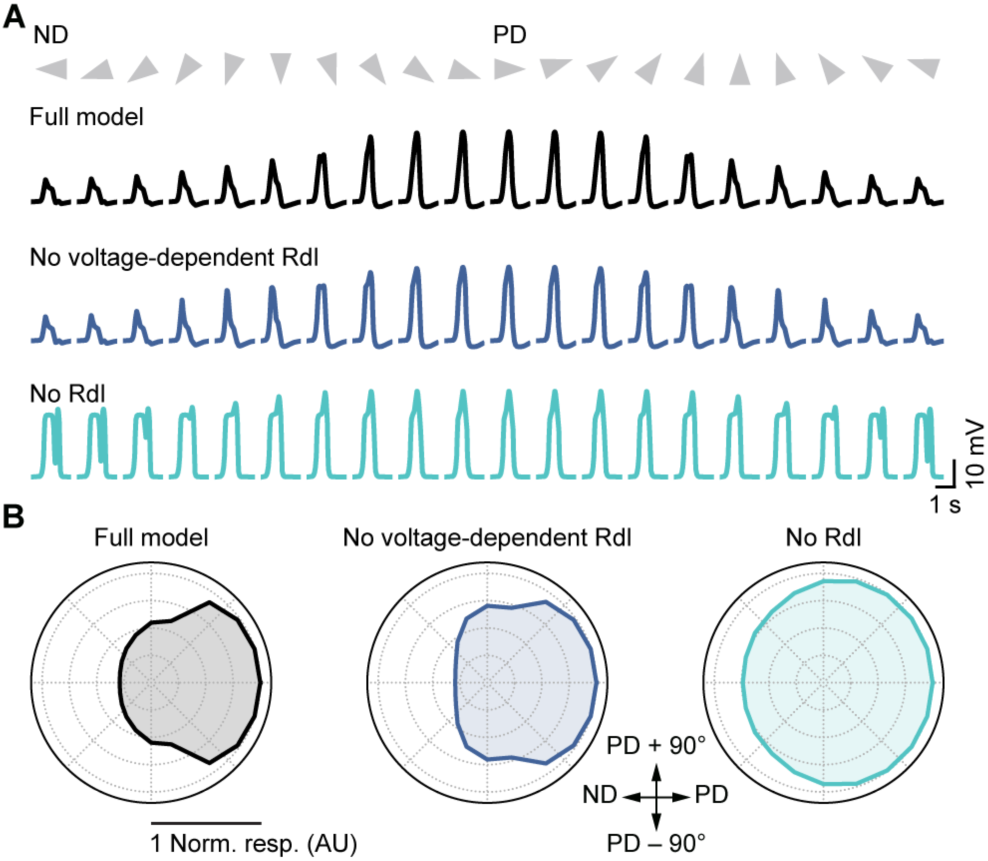
A model of how voltage-dependent inhibition sharpens directional tuning. (A) A biophysically realistic, active T4 neuron model was stimulated with sequences of synaptic input mimicking motion in 20 directions (arrowheads). The full model (black) incorporated ligand- and voltage- gated properties of Rdl and produced sharply tuned membrane potential responses at the axon terminal (black). Blocking of the voltage-gated component of Rdl broadened the tuning (blue), and complete removal of Rdl caused a drastic reduction in response selectivity (cyan). (B) Simulations from (A) represented as polar plots of peak-aligned mean responses normalized to the PD-response of the respective model. PD, preferred direction; ND, null direction; AU, arbitrary units. See also STAR Methods.

### Targeted Rdl receptor knockdown disrupts behavior

Seeing that interference with *Rdl* in T4 and T5 neurons impairs directional tuning (Figures 3F, 3G, 4, and S7), we asked whether this cell type-specific functional change also affects the fly at a larger scale. Therefore, we tested visuomotor behavior of the flies in open- and closed-loop settings using tethered fly-on-ball walking assays. In open loop, control flies responded to a horizontally moving (60° s^−1^) contrast edge with a typical optomotor response that consisted of a turn in the direction of visual motion. The rotational velocity was lower when the stimulus moved in the back-to-front direction compared to the front-to-back direction (Figure 5A).^57^ Flies with *Rdl* RNAi targeted to T4 and T5 neurons still turned in the expected direction, which demonstrates that they are not motion blind, but the distinction in turning amplitude between back-to-front and front-to-back motion disappeared completely (Figure 5A). The characteristic response difference between back-to-front and front-to-back motion in control flies is well in line with their behavioral tendency to maintain a dark vertical bar in center view under closed-loop conditions.^20,57^ The lack of response difference in flies with *Rdl*-deficient T4 and T5 neurons (Figure 5A) consequently suggests that these flies should lose this ability. Further experiments confirmed the prediction: Flies with T4/T5-specific knockdown of *Rdl* not only stopped fixating the 10°-wide bar in front, but they actively avoided the bar by turning away from it (Figure 5B). This complete inversion of behavioral preference due to knockdown of a single type of GABA_A r_eceptor in a genetically defined population of neurons speaks to the importance of precisely timed and balanced inhibition.

**Figure 5.**
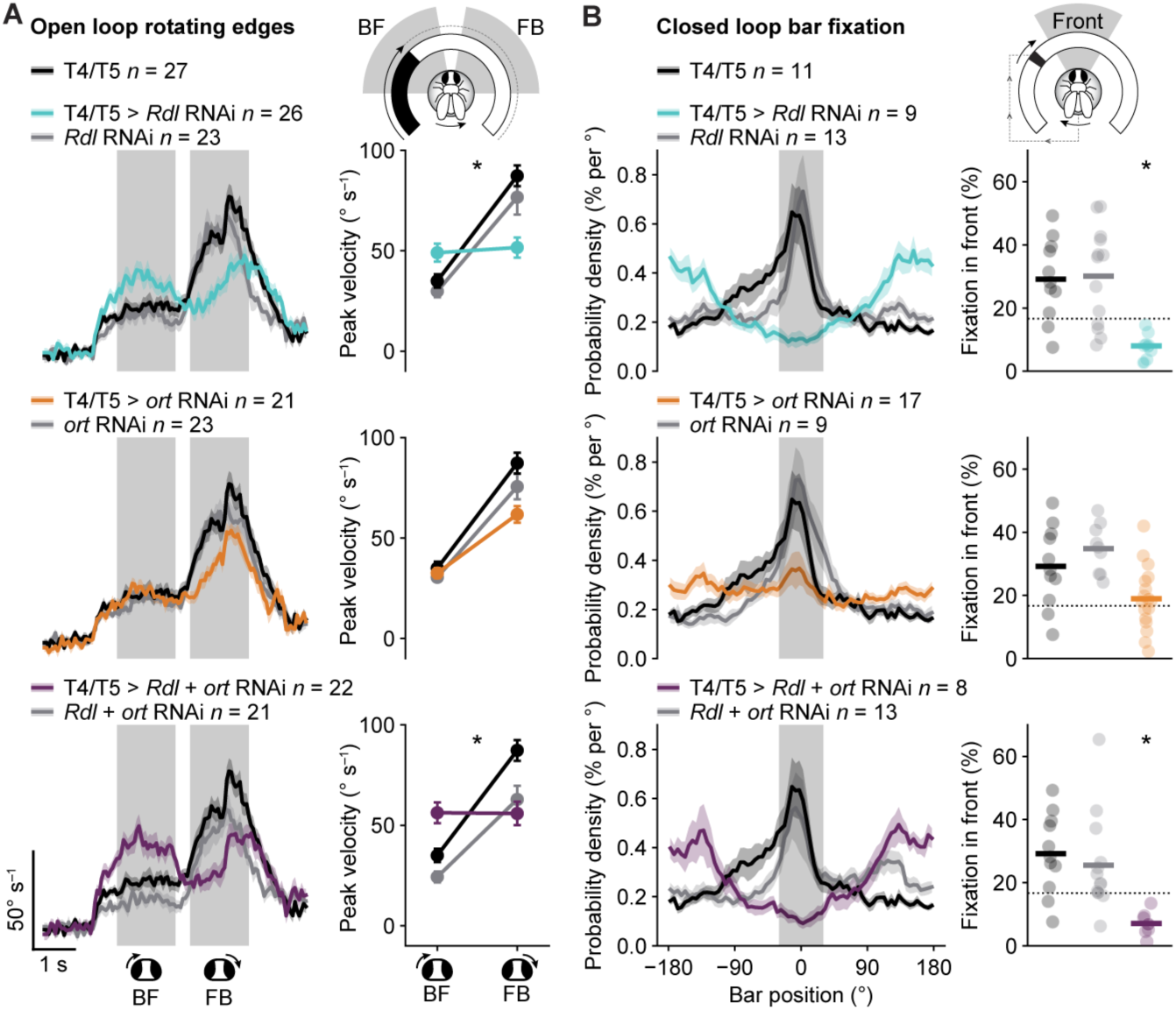
T4/T5-targeted Rdl knockdown changes visuomotor behavior. (A) Rotational velocity of flies expressing *Rdl* RNAi (cyan, top), *ort* RNAi (orange, center), or *Rdl* + *ort* RNAi (purple, bottom) in T4 and T5 neurons and their parental controls (black and gray) in the direction of visual stimulation with single ON or OFF edges moving with 60° s^−1^ along the azimuth around the fly (left) and corresponding peak velocity during time periods (gray shading) of back-to-front (BF) and front-to-back (FB) stimulation (right). Data represent mean ± SEM. Significant differences from parental controls in peak rotational velocity change (FB – BF), determined using Dunn’s multiple comparisons test following Kruskal–Wallis test, are indicated; \**P* = 1.3 × 10^−4^ (T4/T5 > *Rdl* RNAi), \**P* = 1.3 × 10^−3^ (T4/T5 > *Rdl* + *ort* RNAi). (B) Probability density (mean ± SEM) of closed-loop azimuthal bar position (left) and corresponding percentages of time the bar occupied the (gray shaded) frontal 60° window (fixation in front, right). Significant differences from parental controls in fixation in front, determined using Dunn’s multiple comparisons test following Kruskal–Wallis test, are indicated; \**P* = 1.4 × 10^−3^ (T4/T5 > *Rdl* RNAi), \**P* = 9.3 × 10^−3^ (T4/T5 > *Rdl* + *ort* RNAi). Light circles indicate individual flies; horizontal lines indicate the genotype averages; dotted lines indicate chance level. Driver control data (T4/T5, black) are repeated in each panel for comparison. *n* values indicate the number of flies. See also Tables S1 and S3, and STAR Methods.

## Discussion

Connectomes supply wiring diagrams of neural circuits; transcriptomes catalogue their molecular parts. Yet, as of now, neither connectivity matrices nor mRNA inventories predict how a neuron computes, because single-neuron computations are tied to the intrinsic biophysical properties of receptors and channels.^56^ Here, guided by connectomes^12–17^ and transcriptomes,^24–27^ we identify the molecular components of a neural computation^58^ and characterize them biophysically.

In the dendrites of the elementary motion detector neurons T4 and T5, the computation of visual motion has been proposed to rely on well-timed, shunting inhibition that suppresses activity whenever a visual stimulus moves in a non-preferred direction.^2,8–11^ We characterize the neurotransmitter receptors present at inhibitory synapses (Figures 2D–2G) and find that they recruit both the GABA_A r_eceptor Rdl and the histamine receptor Ort, which process GABA signals with different ligand sensitivities and temporal dynamics (Figures 2H–2L). Why Ort—a histamine-gated chloride channel involved in photoreception—should also reside at GABAergic synapses poses an intriguing evolutionary puzzle. Although our experiments failed to assign a clear developmental or physiological function to the spatially confined presence of Ort on the dendrites of T4 and T5 neurons, its sensitivity to high concentrations of GABA hints at a dynamic reserve.

Even though absolute transmitter concentrations in the synaptic cleft remain unknown, it is possible that, under specific lighting or stimulus conditions, the local GABA concentration exceeds the sensitivity of Rdl. In such a high-concentration regime, Ort could step in to rescue GABA responses. Targeting two receptors with shared ligand specificity but distinct sensitivities to the same synapse may thus provide a means of extending the system’s dynamic range and retaining motion vision under extreme conditions. Another open question concerns the stoichiometry of the multi-subunit cys-loop receptors and the possibility of heteromeric assemblies. Although co-localization of the two receptor subunits and the absence of histamine responses in S2R+ cells co-expressing them suggest that Rdl and Ort form heteromeric receptors, this awaits verification by, for example, co-immunoprecipitation.

The main weight of null-direction suppression is carried by Rdl (Figures 3F, 3G, and S7), which we find to be anything but a passive GABA-gated pore. Rdl exhibits ligand-independent, intrinsic sensitivity to membrane potential (Figures 1H and 1I). Previous studies have reported similar subunit-dependent outward rectification of other GABA_A r_eceptor currents,^59–62^ which, however, are contingent on agonist binding. The presence of two completely distinct gating modes in one ion channel has been described before, for example, for KCNQ2 potassium channels,^63,64^ and could be more widespread than the conventional categories of ligand- and voltage-gated channels imply. In this case, the voltage-sensitivity of Rdl might have gone unnoticed up until now due to baseline subtraction routines that focused on transmitter-induced currents.^65–67^ Our simulations suggest that ligand- and voltage-gating of Rdl have distinct roles in computing visual motion (Figure 4). Testing them experimentally will depend on the successful identification and perturbation of the protein domains that confer ligand- and voltage-sensitivity. Since cys-loop ligand-gated ion channels are structurally and phylogenetically related to voltage-gated ion channels,^68^ a voltage-sensing domain of Rdl might be more than a vestigial feature and still serve its archetypal purpose.

Genetic knockdown of Rdl in T4 and T5 neurons breaks their ability to compute visual motion: Rdl-deficient neurons respond indiscriminately to non-motion stimuli and their directional tuning worsens drastically (Figures 3E–3G and S7), to the extent that it affects optomotor responses and changes closed-loop fixation behavior completely (Figure 5). Computing and perceiving visual motion, it turns out, depends not only on synaptic wiring, but also on how the complex gating modes of ionotropic receptors format and interpret synaptic signals. The GABA sensitivity of Ort and the voltage gating of Rdl expand the molecular repertoire of inhibitory mechanisms in single-neuron computations.

## Resource availability

### Lead contact

Requests for further information and resources should be directed to and will be fulfilled by the lead contact, Birte Zuidinga.

### Materials availability

Plasmids generated in this study have been deposited to Addgene and will be available as of the date of final publication. Fly lines generated in this study are available from the lead contact upon request.

### Data and code availability

Confocal microscopy, electrophysiology, two-photon imaging, preprocessed RNA sequencing^24^ and behavioral data have been deposited at Zenodo and will be publicly available as of the date of final publication. This paper analyzes existing, publicly available RNA sequencing data, accessible at Gene Expression Omnibus (GEO: GSE147987, GEO: GSE156455 accessed from https://doi.org/10.5281/zenodo.17932486, GEO: GSE142787, and GEO: GSE116969).

This paper analyzes existing microscopy data on the subcellular localization of GluClα, Dα7, and Rdl provided by R.M.V. that were published before.^22^ This paper analyzes existing, publicly available dendrite morphology and neurotransmitter prediction data from the FlyWire FAFB v783 connectome,^14–17^ accessible at codex.flywire.ai (cell IDs 720575940612221669 and 720575940627480842).

Original code to reproduce analyses, figures, and computational models has been deposited at Github and will be publicly available as of the date of final publication.

Any additional information required to reanalyze the data reported in this paper is available from the lead contact upon request.

## Acknowledgments

We thank Ronald Davis, Barry Dickson, Sandra Fendl, Chun Han, Robin Hiesinger, Chi-Hon Lee, Yulong Li, Aljoscha Nern, Gerald M. Rubin, Tabea Schilling, Takashi Suzuki, the Janelia Research Campus and the TRiP at Harvard Medical School (NIH/NIGMS R01-GM084947) for flies; the Drosophila Genomics Resource Center (NIH 2P40OD010949) for S2R+ cells; Thomas Kornberg for the *pAc5>GAL4* plasmid, Gerald M. Rubin for the *pJFRC7-20XUAS-IVS-mCD8::GFP* plasmid, and Rosa Barrio & James D. Sutherland for the *pAc5-STABLE2-neo* plasmid; Ronald Davis for rabbit anti-Rdl and Lawrence Zipursky for rat anti-Ort antibody; René Rost for assistance with cell culture; Michael Drews and Aljoscha Leonhardt for software; Nikolai Hoermann for preprocessed RNA sequencing data, Stefan Prech, Florian Richter, and Jürgen Haag for technical assistance; and Sandra Fendl, Jonatan Malis, Tabea Schilling, and Polina Arzhevikina for preliminary experiments. Flies obtained from the Bloomington Drosophila Stock Center (NIH P40OD018537) and Vienna Drosophila Resource Center were used in this study.

This work was supported by the Max Planck Society and the Medical University of Graz, as well as by the European Research Council Starting Grant 101116996 (TemProDroMe) to L.N.G. Views and opinions expressed are those of the authors only and do not necessarily reflect those of the European Union or the European Research Council Executive Agency.

## Author contributions

Conceptualization, B.Z., J.P.M., A.B., and L.N.G.; methodology, B.Z., J.P.M., R.M.V., E.S., C.C., M.Ö., and L.N.G.; investigation, B.Z., J.P.M., R.M.V., E.S., C.C., and L.N.G.; writing – original draft, B.Z.; writing – review & editing, J.P.M., R.M.V., E.S., C.C., M.Ö., A.B., and L.N.G.; funding acquisition, A.B. and L.N.G.; resources, B.Z., J.P.M., R.M.V., E.S., C.C., M.Ö., and L.N.G.; supervision, J.P.M., A.B., and L.N.G.

## Declaration of interests

The authors declare no competing interests.

## Supplemental information

Document S1. Figures S1–S8 Document S2. Tables S1–S3

## STAR★METHODS

### Key resources table

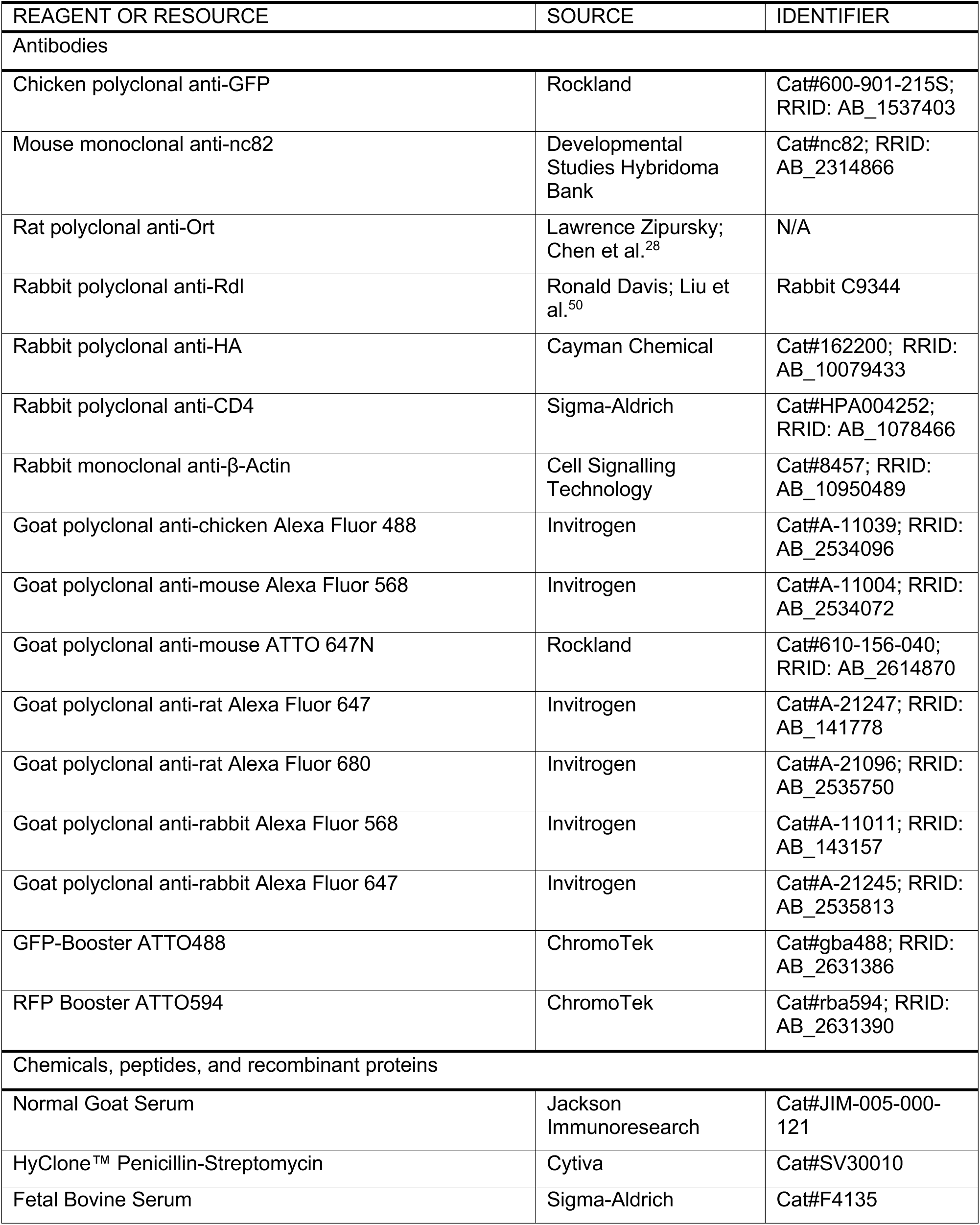
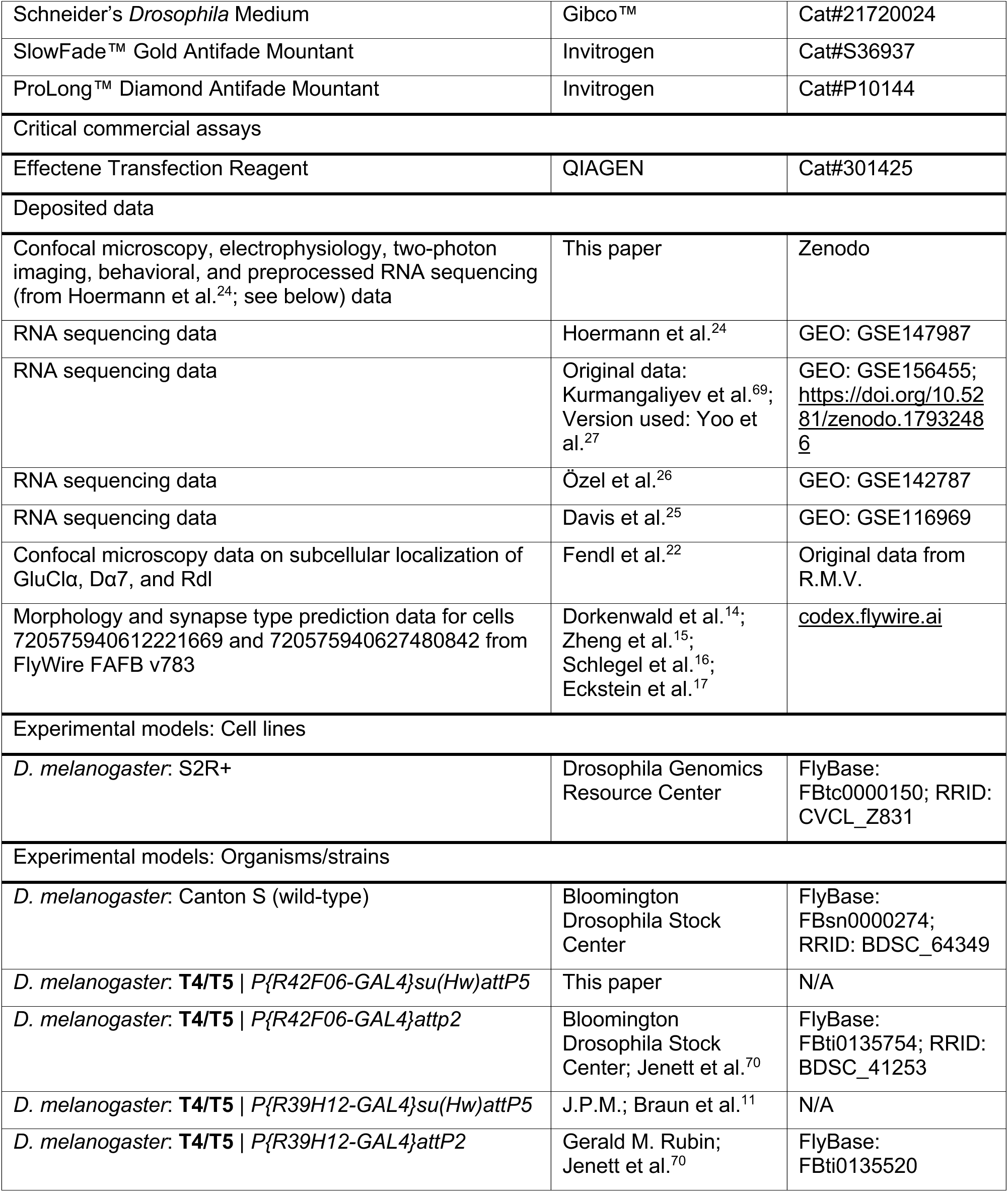
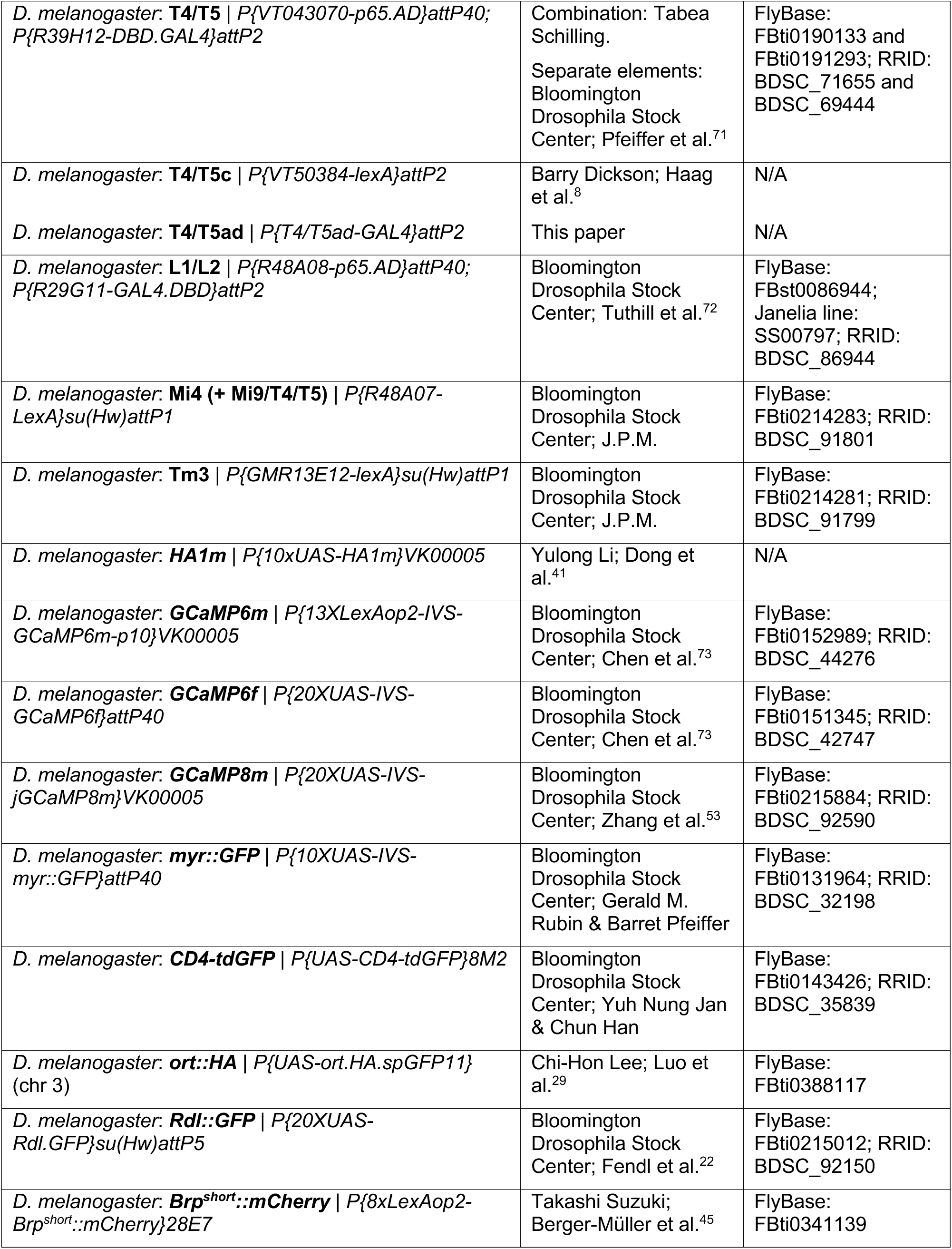
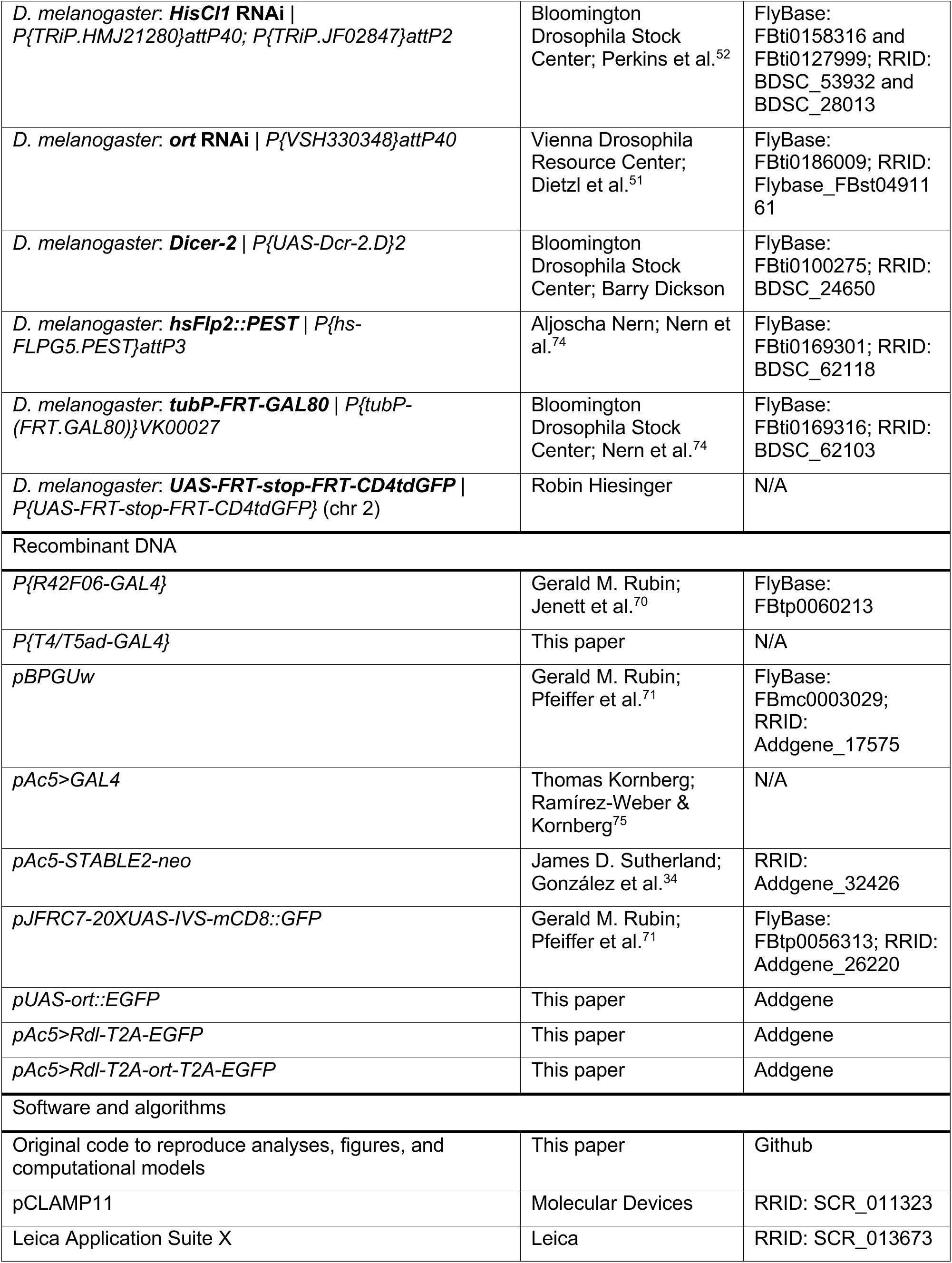
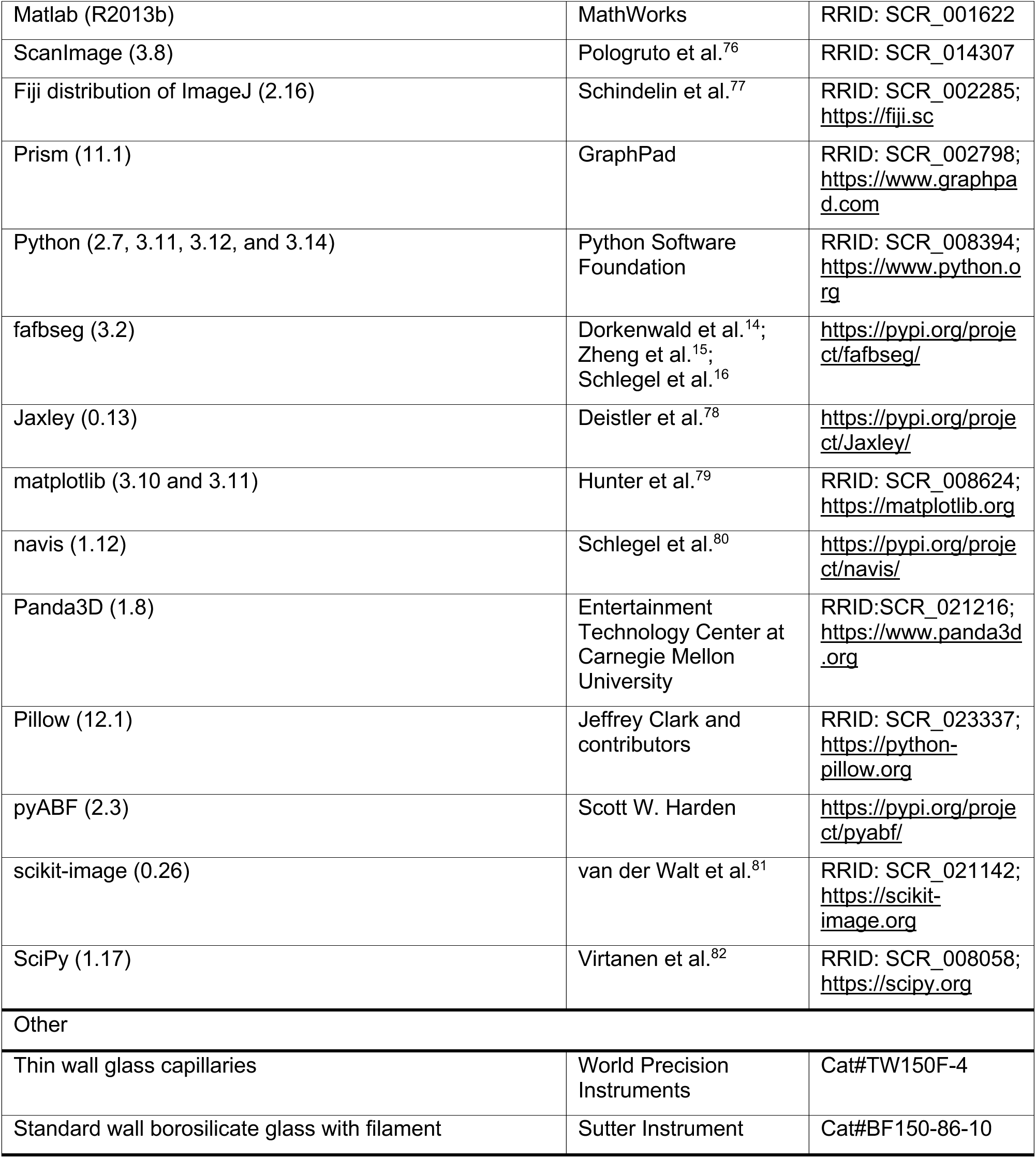

| REAGENT OR RESOURCE | SOURCE | IDENTIFIER |
|---|---|---|
| Antibodies |  |  |
| Chicken polyclonal anti-GFP | Rockland | Cat#600-901-215S; RRID: AB_1537403 |
| Mouse monoclonal anti-nc82 | Developmental Studies Hybridoma Bank | Cat#nc82; RRID: AB_2314866 |
| Rat polyclonal anti-Ort | Lawrence Zipursky; Chen et al.<sup>28</sup> | N/A |
| Rabbit polyclonal anti-Rdl | Ronald Davis; Liu et al.<sup>50</sup> | Rabbit C9344 |
| Rabbit polyclonal anti-HA | Cayman Chemical | Cat#162200; RRID: AB_10079433 |
| Rabbit polyclonal anti-CD4 | Sigma-Aldrich | Cat#HPA004252; RRID: AB_1078466 |
| Rabbit monoclonal anti-β-Actin | Cell Signalling Technology | Cat#8457; RRID: AB_10950489 |
| Goat polyclonal anti-chicken Alexa Fluor 488 | Invitrogen | Cat#A-11039; RRID: AB_2534096 |
| Goat polyclonal anti-mouse Alexa Fluor 568 | Invitrogen | Cat#A-11004; RRID: AB_2534072 |
| Goat polyclonal anti-mouse ATTO 647N | Rockland | Cat#610-156-040; RRID: AB_2614870 |
| Goat polyclonal anti-rat Alexa Fluor 647 | Invitrogen | Cat#A-21247; RRID: AB_141778 |
| Goat polyclonal anti-rat Alexa Fluor 680 | Invitrogen | Cat#A-21096; RRID: AB_2535750 |
| Goat polyclonal anti-rabbit Alexa Fluor 568 | Invitrogen | Cat#A-11011; RRID: AB_143157 |
| Goat polyclonal anti-rabbit Alexa Fluor 647 | Invitrogen | Cat#A-21245; RRID: AB_2535813 |
| GFP-Booster ATTO488 | ChromoTek | Cat#gba488; RRID: AB_2631386 |
| RFP Booster ATTO594 | ChromoTek | Cat#rba594; RRID: AB_2631390 |
| Chemicals, peptides, and recombinant proteins |  |  |
| Normal Goat Serum | Jackson Immunoresearch | Cat#JIM-005-000-121 |
| HyClone™ Penicillin-Streptomycin | Cytiva | Cat#SV30010 |
| Fetal Bovine Serum | Sigma-Aldrich | Cat#F4135 |
| Schneider’s *Drosophila* Medium | Gibco™ | Cat#21720024 |
| SlowFade™ Gold Antifade Mountant | Invitrogen | Cat#S36937 |
| ProLong™ Diamond Antifade Mountant | Invitrogen | Cat#P10144 |
| Critical commercial assays |  |  |
| Effectene Transfection Reagent | QIAGEN | Cat#301425 |
| Deposited data |  |  |
| Confocal microscopy, electrophysiology, two-photon imaging, behavioral, and preprocessed RNA sequencing (from Hoermann et al.<sup>24</sup>; see below) data | This paper | Zenodo |
| RNA sequencing data | Hoermann et al.<sup>24</sup> | GEO: GSE147987 |
| RNA sequencing data | Original data: Kurmangaliyev et al.<sup>69</sup>; Version used: Yoo et al.<sup>27</sup> | GEO: GSE156455; https://doi.org/10.5281/zenodo.17932486 |
| RNA sequencing data | Özel et al.<sup>26</sup> | GEO: GSE142787 |
| RNA sequencing data | Davis et al.<sup>25</sup> | GEO: GSE116969 |
| Confocal microscopy data on subcellular localization of GluClα, Dα7, and Rdl | Fendl et al.<sup>22</sup> | Original data from R.M.V. |
| Morphology and synapse type prediction data for cells 720575940612221669 and 720575940627480842 from FlyWire FAFB v783 | Dorkenwald et al.<sup>14</sup>; Zheng et al.<sup>15</sup>; Schlegel et al.<sup>16</sup>; Eckstein et al.<sup>17</sup> | codex.flywire.ai |
| Experimental models: Cell lines |  |  |
| *D. melanogaster*: S2R+ | Drosophila Genomics Resource Center | FlyBase: FBtc0000150; RRID: CVCL_Z831 |
| Experimental models: Organisms/strains |  |  |
| *D. melanogaster*: Canton S (wild-type) | Bloomington Drosophila Stock Center | FlyBase: FBsn0000274; RRID: BDSC_64349 |
| *D. melanogaster*: **T4/T5** \| *P{R42F06-GAL4}su(Hw)attP5* | This paper | N/A |
| *D. melanogaster*: **T4/T5** \| *P{R42F06-GAL4}attp2* | Bloomington Drosophila Stock Center; Jenett et al.<sup>70</sup> | FlyBase: FBti0135754; RRID: BDSC_41253 |
| *D. melanogaster*: **T4/T5** \| *P{R39H12-GAL4}su(Hw)attP5* | J.P.M.; Braun et al.<sup>11</sup> | N/A |
| *D. melanogaster*: **T4/T5** \| *P{R39H12-GAL4}attP2* | Gerald M. Rubin; Jenett et al.<sup>70</sup> | FlyBase: FBti0135520 |
| *D. melanogaster*: **T4/T5** \| *P{VT043070-p65.AD}attP40; P{R39H12-DBD.GAL4}attP2* | Combination: Tabea Schilling.<br>Separate elements: Bloomington Drosophila Stock Center; Pfeiffer et al.<sup>71</sup> | FlyBase: FBti0190133 and FBti0191293; RRID: BDSC_71655 and BDSC_69444 |
| *D. melanogaster*: **T4/T5c** \| *P{VT50384-lexA}attP2* | Barry Dickson; Haag et al.<sup>8</sup> | N/A |
| *D. melanogaster*: **T4/T5ad** \| *P{T4/T5ad-GAL4}attP2* | This paper | N/A |
| *D. melanogaster*: **L1/L2** \| *P{R48A08-p65.AD}attP40; P{R29G11-GAL4.DBD}attP2* | Bloomington Drosophila Stock Center; Tuthill et al.<sup>72</sup> | FlyBase: FBst0086944; Janelia line: SS00797; RRID: BDSC_86944 |
| *D. melanogaster*: **Mi4 (+ Mi9/T4/T5)** \| *P{R48A07-LexA}su(Hw)attP1* | Bloomington Drosophila Stock Center; J.P.M. | FlyBase: FBti0214283; RRID: BDSC_91801 |
| *D. melanogaster*: **Tm3** \| *P{GMR13E12-lexA}su(Hw)attP1* | Bloomington Drosophila Stock Center; J.P.M. | FlyBase: FBti0214281; RRID: BDSC_91799 |
| *D. melanogaster*: ***HA1m*** \| *P{10xUAS-HA1m}VK00005* | Yulong Li; Dong et al.<sup>41</sup> | N/A |
| *D. melanogaster*: ***GCaMP6m*** \| *P{13XLexAop2-IVS-GCaMP6m-p10}VK00005* | Bloomington Drosophila Stock Center; Chen et al.<sup>73</sup> | FlyBase: FBti0152989; RRID: BDSC_44276 |
| *D. melanogaster*: ***GCaMP6f*** \| *P{20XUAS-IVS-GCaMP6f}attP40* | Bloomington Drosophila Stock Center; Chen et al.<sup>73</sup> | FlyBase: FBti0151345; RRID: BDSC_42747 |
| *D. melanogaster*: ***GCaMP8m*** \| *P{20XUAS-IVS-jGCaMP8m}VK00005* | Bloomington Drosophila Stock Center; Zhang et al.<sup>53</sup> | FlyBase: FBti0215884; RRID: BDSC_92590 |
| *D. melanogaster*: ***myr::GFP*** \| *P{10XUAS-IVS-myr::GFP}attP40* | Bloomington Drosophila Stock Center; Gerald M. Rubin & Barret Pfeiffer | FlyBase: FBti0131964; RRID: BDSC_32198 |
| *D. melanogaster*: ***CD4-tdGFP*** \| *P{UAS-CD4-tdGFP}8M2* | Bloomington Drosophila Stock Center; Yuh Nung Jan & Chun Han | FlyBase: FBti0143426; RRID: BDSC_35839 |
| *D. melanogaster*: ***ort::HA*** \| *P{UAS-ort.HA.spGFP11}* (chr 3) | Chi-Hon Lee; Luo et al.<sup>29</sup> | FlyBase: FBti0388117 |
| *D. melanogaster*: ***Rdl::GFP*** \| *P{20XUAS-Rdl.GFP}su(Hw)attP5* | Bloomington Drosophila Stock Center; Fendl et al.<sup>22</sup> | FlyBase: FBti0215012; RRID: BDSC_92150 |
| *D. melanogaster*: ***Brp<sup>short</sup>::mCherry*** \| *P{8xLexAop2-Brp<sup>short</sup>::mCherry}28E7* | Takashi Suzuki; Berger-Müller et al.<sup>45</sup> | FlyBase: FBti0341139 |
| *D. melanogaster*: ***HisCl1*** **RNAi** \| *P{TRiP.HMJ21280}attP40; P{TRiP.JF02847}attP2* | Bloomington Drosophila Stock Center; Perkins et al.<sup>52</sup> | FlyBase: FBti0158316 and FBti0127999; RRID: BDSC_53932 and BDSC_28013 |
| *D. melanogaster*: ***ort*** **RNAi** \| *P{VSH330348}attP40* | Vienna Drosophila Resource Center; Dietzl et al.<sup>51</sup> | FlyBase: FBti0186009; RRID: Flybase_FBst0491161 |
| *D. melanogaster*: ***Dicer-2*** \| *P{UAS-Dcr-2.D}2* | Bloomington Drosophila Stock Center; Barry Dickson | FlyBase: FBti0100275; RRID: BDSC_24650 |
| *D. melanogaster*: ***hsFlp2::PEST*** \| *P{hs-FLPG5.PEST}attP3* | Aljoscha Nern; Nern et al.<sup>74</sup> | FlyBase: FBti0169301; RRID: BDSC_62118 |
| *D. melanogaster*: ***tubP-FRT-GAL80*** \| *P{tubP-(FRT.GAL80)}VK00027* | Bloomington Drosophila Stock Center; Nern et al.<sup>74</sup> | FlyBase: FBti0169316; RRID: BDSC_62103 |
| *D. melanogaster*: ***UAS-FRT-stop-FRT-CD4tdGFP*** \| *P{UAS-FRT-stop-FRT-CD4tdGFP}* (chr 2) | Robin Hiesinger | N/A |
| Recombinant DNA |  |  |
| *P{R42F06-GAL4}* | Gerald M. Rubin; Jenett et al.<sup>70</sup> | FlyBase: FBtp0060213 |
| *P{T4/T5ad-GAL4}* | This paper | N/A |
| *pBPGUw* | Gerald M. Rubin; Pfeiffer et al.<sup>71</sup> | FlyBase: FBmc0003029; RRID: Addgene_17575 |
| *pAc5>GAL4* | Thomas Kornberg; Ramírez-Weber & Kornberg<sup>75</sup> | N/A |
| *pAc5-STABLE2-neo* | James D. Sutherland; González et al.<sup>34</sup> | RRID: Addgene_32426 |
| *pJFRC7-20XUAS-IVS-mCD8::GFP* | Gerald M. Rubin; Pfeiffer et al.<sup>71</sup> | FlyBase: FBtp0056313; RRID: Addgene_26220 |
| *pUAS-ort::EGFP* | This paper | Addgene |
| *pAc5>Rdl-T2A-EGFP* | This paper | Addgene |
| *pAc5>Rdl-T2A-ort-T2A-EGFP* | This paper | Addgene |
| Software and algorithms |  |  |
| Original code to reproduce analyses, figures, and computational models | This paper | Github |
| pCLAMP11 | Molecular Devices | RRID: SCR_011323 |
| Leica Application Suite X | Leica | RRID: SCR_013673 |
| Matlab (R2013b) | MathWorks | RRID: SCR_001622 |
| ScanImage (3.8) | Pologruto et al.<sup>76</sup> | RRID: SCR_014307 |
| Fiji distribution of ImageJ (2.16) | Schindelin et al.<sup>77</sup> | RRID: SCR_002285; https://fiji.sc |
| Prism (11.1) | GraphPad | RRID: SCR_002798; https://www.graphpad.com |
| Python (2.7, 3.11, 3.12, and 3.14) | Python Software Foundation | RRID: SCR_008394; https://www.python.org |
| fafbseg (3.2) | Dorkenwald et al.<sup>14</sup>; Zheng et al.<sup>15</sup>; Schlegel et al.<sup>16</sup> | https://pypi.org/project/fafbseg/ |
| Jaxley (0.13) | Deistler et al.<sup>78</sup> | https://pypi.org/project/Jaxley/ |
| matplotlib (3.10 and 3.11) | Hunter et al.<sup>79</sup> | RRID: SCR_008624; https://matplotlib.org |
| navis (1.12) | Schlegel et al.<sup>80</sup> | https://pypi.org/project/navis/ |
| Panda3D (1.8) | Entertainment Technology Center at Carnegie Mellon University | RRID:SCR_021216; https://www.panda3d.org |
| Pillow (12.1) | Jeffrey Clark and contributors | RRID: SCR_023337; https://python-pillow.org |
| pyABF (2.3) | Scott W. Harden | https://pypi.org/project/pyabf/ |
| scikit-image (0.26) | van der Walt et al.<sup>81</sup> | RRID: SCR_021142; https://scikit-image.org |
| SciPy (1.17) | Virtanen et al.<sup>82</sup> | RRID: SCR_008058; https://scipy.org |
| Other |  |  |
| Thin wall glass capillaries | World Precision Instruments | Cat#TW150F-4 |
| Standard wall borosilicate glass with filament | Sutter Instrument | Cat#BF150-86-10 |

## Experimental model details

### Flies

*Drosophila melanogaster* flies were cultivated on a cornmeal, molasses and yeast medium under a 12–12 hours light–dark cycle at 25 °C and 60% humidity. All of the experiments were carried out using female flies, except for immunohistochemical experiments to determine co-localization of receptor proteins with presynaptic terminals (Figures 2D–2G), for which flies of both sexes were used. Genotypes of all experimental flies can be found in Table S3; the details of separate transgenes are listed in the Key Resources Table. For generating the *R42F06-GAL4* (T4/T5) driver line on the second chromosome, the *P{R42F06-GAL4}* plasmid^70^ was injected into the *su(Hw)attP5* landing site (RRID: BDSC_34765) for PhiC31 integrase-mediated transgenesis by BestGene, Inc. For generating the T4/T5ad*-GAL4* driver line, the enhancer sequence (chr2L: 12,757,364–12,757,950) was chosen based on ATACseq data indicating it to be a highly accessible genomic region in T4 neurons.^83^ The enhancer sequence was synthesized by Genewiz (Azenta) and inserted into a *pBPGUw* plasmid backbone^71^ digested with ZraI and Nael. The plasmid was injected into the *attP2* landing site (RRID: BDSC_8622) for PhiC31 integrase-mediated transgenesis by BestGene, Inc. No ethical approval was required for research on *Drosophila melanogaster*.

### Cells

The *Drosophila melanogaster* S2R+ cell line (RRID: CVCL_Z831),^33^ originating from late embryonic stage male Oregon R cells, was obtained from the Drosophila Genomics Resource Center (Bloomington, IN). Cells were cultured at 25 °C in Schneider’s *Drosophila* medium supplemented with 10% fetal bovine serum (Sigma-Aldrich, Cat#F4135) and penicillin/streptomycin (Cytiva, Cat#SV30010). Cultures were passaged every 3–4 days. For electrophysiological experiments, cells were plated on round glass coverslips (25 mm diameter, VWR) in 6-well plates (Falcon). One day after plating, cells at approximately 40% confluency were transfected using the Effectene Transfection Reagent (QIAGEN, Cat#301425). Cells were transfected with 0.25 μg per well of each *pAc5>GAL4* and *pUAS-ort::EGFP* (*ort*), or 0.5 μg plasmid DNA per well of *pAc5-STABLE2-neo* (control), *pAc5>Rdl-T2A-EGFP* (*Rdl*), or *pAc5>Rdl-T2A-ort-T2A-EGFP* (*Rdl* + *ort*).

## Method details

### Plasmids

Five different plasmids were used to express receptor proteins and fluorophores in S2R+ cells. The *pAc5>GAL4* plasmid, driving *GAL4* expression under the *Actin* promotor, was a gift from Thomas Kornberg.^75^ The *pUAS-ort::EGFP* plasmid, expressing *ort* labelled with *EGFP* under control of *GAL4*, was generated as previously described^22^ by synthesizing the coding DNA sequence (CDS) of *ort* with an *EGFP* insertion (with a flexible 4×GGS linker) between exons 3 and 4. The sequence was flanked with NotI and XbaI restriction sites to be inserted into the *pJFRC7-20XUAS-IVS-mCD8::GFP* plasmid (RRID: Addgene_26220)^70^ using restriction cloning. The *pAc5-STABLE2-neo* plasmid (RRID: Addgene_32426)^34^ expressing *mCherry* and *EGFP* under the *Actin* promoter was used for the control condition to transfect cells without receptor expression and served as backbone to generate the *pAc5>Rdl-T2A-EGFP* (*Rdl*) and *pAc5>Rdl-T2A-ort-T2A-EGFP* (*Rdl* + *ort)* plasmids. For these plasmids, synthesized *Rdl* CDS (isoform D) was cloned into the backbone alone or T2A-linked with *ort* CDS, using the EcoRI and NotI restriction sites. DNA synthesis and restriction cloning were carried out by Genewiz (Azenta).

### Patch-clamp electrophysiology

Cells were used for electrophysiological recordings from 2 until 4 days after transfection. One coverslip containing cells was placed inside the lid of a 35-mm cell culture dish (Falcon) and submerged in extracellular solution containing 5 mM TES, 103 mM NaCl, 3 mM KCl, 26 mM NaHCO_3,_ 1 mM NaH_2P_O_4,_ 1.5 mM CaCl_2,_ 4 mM MgCl_2,_ 10 mM trehalose, 10 mM glucose and 7 mM sucrose (pH 7.3, 280 mOsM, equilibrated with 5% CO_2 a_nd 95% O_2)_. For experiments with a reduced extracellular chloride concentration, NaCl was substituted by sodium aspartate, resulting in a total chloride concentration of 14 mM (low) instead of 117 mM (normal). Cells were visualized using an InVivo SliceScope (Scientifica) microscope equipped with a ×60/1.0 NA water-immersion objective (LUMPLFLN60XW, Olympus) and white LED transmitted light illumination. Transfected cells were identified by their GFP signal (Figure 1G) using epifluorescence microscopy with an LQ-HXP 120 light source (Leistungselektronik Jena). Micropipettes were pulled from borosilicate glass capillaries with an outer diameter of 1.5 mm and an inner diameter of 1.12 mm (World Precision Instruments, Cat#TW150F-4) or 0.86 mm (Sutter Instrument, Cat#BF150-86-10) using a P-97 Flaming/Brown (Sutter Instrument) or PC-100 (Narishige) puller, respectively. Recording pipettes were filled with intracellular solution containing 10 mM HEPES, 140 mM potassium aspartate, 1 mM KCl, 4 mM MgATP, 0.5 mM Na_3G_TP and 1 mM EGTA (pH 7.3, 265 mOsM). For neurotransmitter applications, pipettes were filled with extracellular solution containing either 1 mM or 50 mM GABA, histamine, or glutamate, and coupled to a PDES-02DX pneumatic drug ejection system (NPI Electronic) that delivered 50 kPa pressure pulses when triggered. After establishing whole-cell voltage-clamp configuration and positioning the transmitter application pipette within ∼20 µm of the cell, the stimulus protocol was started.

The stimulus consisted of 17 sweeps of 5 s each, of different clamp potentials (−120 mV to +40 mV, with 10 mV increments) lasting 3.5 s, with a baseline clamp potential of −70 mV before and after (Figure 1G). A 20-ms pulse triggered neurotransmitter ejection 1.05 s after the change in clamp potential. Stimuli were repeated at least three times and averaged online, except when the recording became unstable and was stopped prematurely, which resulted in one or two averaged repetitions. Signals were recorded using a MultiClamp 700B amplifier (Molecular Devices), sampled at 10 kHz and low-pass-filtered using a Digidata 1550B digitizer controlled by pCLAMP11 software (Molecular Devices). Clamp potentials were corrected for the liquid junction potentials corresponding to the respective solutions, which were calculated to be −16 mV (normal [Cl^−^]_out)_ and −9 mV (low [Cl^−^]_out)_. Experiments were done at room temperature (21–23 °C). Data were imported in Python v.3.12 (Python Software Foundation) using the pyABF package (v.2.3) and were analyzed using custom-written routines. The exemplary brightfield image (Figure 1G) was taken after establishing whole-cell configuration with an *Rdl*-expressing cell. Native EGFP signal was gaussian smoothed (σ = 5 pixels) and overlaid.

To quantify transmitter-elicited current responses, the measured currents were baseline subtracted using the average current during the 220 ms prior to the transmitter application trigger. Peak responses were defined as the current at the time point of the maximal standard deviation between all traces (corresponding to the clamp potentials) of a unique cell, to prevent trigger artefacts from being mistaken as responses. The decay of the response to 50 mM GABA was quantified at +40 mV clamp potential by fitting an exponential decay function to the normalized current starting at 80% of its peak value using optimize.curve_fit from the SciPy (v.1.17) package.^82^ Cells whose current decay could not be fit adequately within the 2 s after the peak (4/13 of *Rdl*- and 1/11 of *ort*-expressing cells) were excluded from the analysis.

Reversal potentials were calculated from measurements of responding cells (peak standard deviation > 0.1 nA; 13/15 of *Rdl*-expressing cells at normal [Cl^−^]_out a_nd all recorded cells in the other conditions) by linear interpolation of the current-voltage values before and after the zero-crossing. Voltage-elicited currents were quantified in recordings without transmitter-related responses (the 50 mM glutamate condition for receptor-expressing cells and the 50 mM GABA condition for control cells) by subtracting the average baseline current in the 175 ms preceding the voltage step from the average steady-state current following the voltage step, during the 220 ms period before the trigger of neurotransmitter application.

### Immunofluorescence

Primary antibodies (and corresponding concentrations) used were chicken anti-GFP (1:500, Rockland, Cat#600901215S, RRID: AB_1537403), mouse anti-nc82 (1:20, DSHB, Cat#nc82, RRID: AB_2314866; to stain Bruchpilot, or Brp), rat anti-Ort (1:200, gift from Lawrence Zipursky),^28^ rabbit anti-Rdl (1:100, gift from Ronald Davis, C9344),^50^ rabbit anti-HA (1:300, Cayman Chemical, Cat#162200, RRID: AB_10079433), rabbit anti-CD4 (1:200, Sigma-Aldrich, Cat#HPA004252, RRID: AB_1078466), and rabbit anti-β-Actin (1:200, Cell Signalling Technology, Cat#8457, RRID: AB_10950489). Secondary antibodies (and corresponding concentrations) used were goat anti-chicken Alexa Fluor 488 (1:500, Invitrogen, Cat#A-11039, RRID: AB_2534096), goat anti-mouse Alexa Fluor 568 (1:500, Invitrogen, Cat#A-11004, RRID: AB_2534072), goat anti-mouse ATTO 647N (1:300, Rockland, Cat#610-156-040, RRID: AB_2614870), goat anti-rat Alexa Fluor 647 (1:500, Invitrogen, Cat#A-21247, RRID: AB_141778), goat anti-rat Alexa Fluor 680 (1:400, Invitrogen, Cat#A-21096, RRID: AB_2535750), goat anti-rabbit Alexa Fluor 568 (1:400, Invitrogen, Cat#A-11011, RRID: AB_143157), and goat anti-rabbit Alexa Fluor 647 (1:500, Invitrogen, Cat#A-21245, RRID: AB_2535813). Fluorescence boosters (and corresponding concentrations) used were GFP-Booster ATTO 488 (1:400, Cat#gba488, RRID: AB_2631386) and RFP-Booster ATTO 594 (1:200, Cat#rba594, RRID: AB_2631390), both from ChromoTek.

Fly brains (10 hours–6 days old) of the genotypes of interest were dissected in phosphate buffered saline (PBS; 137 mM NaCl, 3 mM KCl, 8 mM Na_2H_PO_4,_ 1.5 mM KH_2P_O_4,_ pH 7.3) and fixed for 20–30 minutes in 4% (w/v) paraformaldehyde in PBS with 0.1% (v/v) Triton X-100. Brains were washed in PBS with 0.3% (v/v) Triton X-100 (PBT) and blocked in PBT containing 10% (v/v) normal goat serum (NGS; Jackson Immunoresearch, Cat#JIM-005-000-121) for one hour at room temperature (21–23 °C) or overnight at 4 °C. After blocking, brains were incubated with primary antibodies diluted in PBT containing 5% (v/v) NGS for at least 48 hours at 4 °C, washed in PBT, and subsequently incubated with secondary antibodies diluted in PBT containing 5% (v/v) NGS for at least 48 hours at 4 °C. After washing in PBT again, brains were mounted in SlowFade™ Gold Antifade Mountant (Invitrogen, Cat#S36937).

Cells for immunofluorescence were prepared and transfected in the same way as cells used for electrophysiology. The staining procedure was carried out at room temperature (21–23 °C). Four days after transfection, cell culture medium was removed from the wells and cells were fixed in 4% (w/v) paraformaldehyde in PBS with 0.1% (v/v) Triton-X for 20 minutes. Cells were washed 3× for 5 minutes in PBT and blocked for 40 minutes in 2% (v/v) NGS in PBS with 0.1% (v/v) Triton-X (blocking buffer). After blocking, cells were incubated with primary antibodies diluted in blocking buffer for 1.5 hours, washed 3× for 5 minutes in PBT, and incubated for one hour in secondary antibodies diluted in blocking buffer. After 3 more washes, coverslips were mounted upside down on microscopy slides in ProLong™ Diamond Antifade Mountant (Invitrogen, Cat#P10144) and cured overnight. All coverslips were stained with chicken anti-GFP and goat anti-chicken Alexa Fluor 488 to visualize transfected cells. Coverslips with cells expressing *pAc5>Rdl-T2A-EGFP* were additionally stained with rabbit anti-Rdl and goat anti-rabbit Alexa 568. Coverslips with cells expressing *pUAS-ort::EGFP* were additionally stained with rat anti-Ort and goat anti-rat Alexa Fluor 680, and rabbit anti-β-Actin and goat anti-rabbit Alexa Fluor 568 (not shown). For cells expressing *pAc5>Rdl-T2A-ort-T2A-EGFP*, antibodies targeting Ort and Rdl were the same as above. Control cells expressing *pAc5-STABLE2-neo* were additionally stained with the same primary and secondary antibodies targeting Ort (not shown), and native mCherry fluorescence was imaged without staining.

Samples were imaged on a Leica Stellaris 5 or Leica TCS SP8 laser scanning confocal microscope (Leica) with HC Plan-apochromat glycerol-immersion objectives (20×/0.75NA and 63×/1.3NA), using the Leica Application Suite X (Leica). Micrographs were processed using the Fĳi distribution of ImageJ.^77^ For proper visualization of fluorescence patterns in exemplary images, contrast and brightness were adjusted separately for each color channel. For quantifications, raw pixel values were used.

### Subcellular Ort localization

To stochastically express *ort::HA* and *myr::GFP* or *CD4::tdGFP* in single T4 or T5 neurons, respectively, a heat shock-based method was used.^22^ Pupae were collected between 4 and 24 hours after puparium formation and heat shocked for 35 minutes at 37 °C. Rabbit anti-HA antibody was used to visualize Ort and chicken anti-GFP antibody to visualize the labelled neurons’ membrane. Background staining of synaptic structures using mouse anti-nc82 antibody facilitated determining dendrite orientations. Single dendrites were imaged in the medulla (T4) and lobula (T5). Maximum intensity z-projections were computed in Fĳi and analyzed further in Python (v.3.12) using Pillow (v.12.1) and SciPy (v.1.17).^82^ Images were cropped to include the whole dendrite and, if necessary, rotated and mirrored to align them to a common frame of reference. Ort fluorescence intensity values were averaged over the distal–proximal axis, and the number of datapoints were equalized to the largest image using linear interpolation. Intensity values were normalized to the maximal value for each image, smoothed with a gaussian filter (σ = 8), and plotted together with averages of previously published data of GluClα, Dα7, and Rdl as reference,^22^ which were reanalyzed the same way.

### Co-localization

Visualization of Ort and Rdl co-localization in T4 and T5 neurons was achieved by simultaneous overexpression of *ort::HA* and *Rdl::GFP*. Co-localization of the receptors with presynaptic neuron terminals was assessed by overexpression of each separate receptor combined with *LexA*/*LexAop*-driven overexpression of *Brp^short^::mCherry* in Mi4 and Tm3 neurons.^45^ The driver line used to target Mi4 neurons (*R48A07-LexA*) targets to a lesser extent also glutamatergic Mi9 and cholinergic T4 and T5 neurons. Ort was fluorescently stained using primary rabbit anti-HA and secondary anti-rabbit Alexa Fluor 647. *Rdl::GFP* and *Brp^short^::mCherry* signals were enhanced using GFP-Booster ATTO 488 and RFP-Booster ATTO 594, respectively. Boosters were added either during the only or, when stained together with Ort, during the second antibody incubation step.

### RNAi validation

The efficacy of *Rdl* and *ort* RNAi to deplete the corresponding receptors was validated by expressing the constructs in all T4 and T5 neurons and visualizing the native proteins in the optic lobe using rabbit anti-Rdl and rat anti-Ort antibody staining, respectively. Control flies expressed *HisCl1* RNAi instead. GCaMP6f (Rdl) or GCaMP6m (Ort) was expressed in addition to mark the neurons and was stained using chicken anti-GFP antibody. Mouse anti-nc82 antibody was used as a counterstain for synaptic structures (Brp). Micrographs were analyzed using custom scripts written in Fiji (ImageJ Macro Language) and Python (v.3.12). Average intensity z-projections of micrographs were made of three central optic lobe slices (z-steps of 0.33 µm), after which the Brp channel was used to delineate the full neuropils (medulla and lobula) manually, and the GFP channel was used to delineate the dendritic regions of T4 and T5 neurons (M10 and Lo1, respectively) using a combined manual and automatic algorithm. The receptor intensity ratio was calculated from the receptor channel by dividing the average intensity of pixels inside the GFP mask by the average intensity of pixels outside the GFP mask within the respective neuropil. Values were normalized to the average values of control flies expressing *HisCl1* RNAi.

### Dendrite morphology

A heat shock-based method^74^ was used to achieve stochastic neuron labelling with *CD4tdGFP* in the absence and presence of *ort* RNAi expression. The used split-*GAL4* driver line targets both T4 and T5 neurons, but brains were mounted with the anterior side facing up to best visualize the shape of T4 dendrites in the medulla. Pupae were collected between 0 and 72 hours after puparium formation and heat shocked for 6 minutes at 37 °C. Membranes were stained using rabbit anti-CD4 antibody. Background staining of synaptic structures using mouse anti-nc82 antibody provided a common frame of reference for the consistent orientation of dendrites. A neuron’s subtype was determined based on the lobula plate layer where its axon terminated. A maximum projection was made of a z-stack incorporating the complete dendrite using Fĳi. Dendritic arbors were rotated such that their longest axis was aligned with the vertical axis of the image. For the automated analysis to work properly, neurons that were visible in the images but not connected to the dendrite of interest were manually removed. Dendrites were only analyzed if they were clearly separable from any adjacent neurons. The morphology of the dendrite was quantified using functions of scikit-image (v.0.26) in Python (v.3.12). Dendrite pixels were defined as pixels with CD4 fluorescence intensity > 90% of the maximum of each image. The dendrite span was determined by taking the convex hull around the dendrite pixels, excluding groups of < 6 connected pixels. The dendrite density was calculated by dividing the number of dendrite pixels in the convex hull by the total number of pixels in the convex hull. The angular orientation of the dendrite pixels in reference to the dendrite origin was visualized for each subtype as a polar histogram using the same method as described before.^24^ The values of each bin represent the percentage of dendrite pixels with an angular orientation within the respective boundaries (using 12 bins of 30°). Using these values as normalized vectors ***v*** corresponding to the (central) angle *ϕ* of each bin, we quantified the dendrite orientation index as the magnitude of the vector sum:

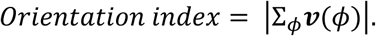

The orientation index can vary between 0 and 1, representing the lowest possible orientation selectivity (the same number of pixels in all bins) and the highest possible orientation selectivity (all pixels in one bin), respectively.

### Two-photon imaging

Microsurgeries and functional imaging were performed using a procedure similar to one described before.^54^ Female flies (1−5 days old) were cold-immobilized and attached to an acrylic post by their thorax, wings, and legs with UV-curing dental glue (Sinfony Opaque Dentin, 3M) solidified using a 440-nm curing light (Woodpecker). The head was tilted forward and inserted into a fitting opening in aluminum foil clamped in a polyoxymethylene recording chamber using beeswax. The posterior side of the head was immersed in the same extracellular solution as used for electrophysiological experiments (normal [Cl^−^]_out)_ without O_2/_CO_2._ Cuticle and soft tissue covering the brain were removed using a hypodermic needle to reveal the right optic lobe. A custom-built two-photon microscope equipped with a 40× water immersion objective (W Plan-Apochromat, NA 1.0, Zeiss) was used to acquire images at 64 × 64 pixels resolution, 12.21 Hz frame rate and 920 nm excitation wavelength, and was controlled by ScanImage software (v.3.8) in Matlab R2013b (MathWorks).^76^ Data were analyzed using custom-written routines in Python (v.2.7 and v.3.12). Motion correction and signal processing were performed using previously described methods.^55^ Three adjacent regions of interest (ROIs) were manually delineated in the central field of view of each recording (to collect signals from only a few overlapping neurons with similar receptive fields), except for calcium imaging experiments using full-field flashes and for histamine imaging, where one larger ROI was delineated to optimize the signal-to-noise ratio. Fluorescence changes (ΔF/F) in each ROI were calculated using an algorithm robust against slow baseline drift.^84^

A custom-built projector-based arena was used to present visual stimuli to the flies.^55^ The arena consists of two micro-projectors (TI DLP Lightcrafter 3000) projecting onto the back of an opaque cylindrical screen covering 180° in azimuth and 105° in elevation of the fly’s visual field. The projectors were programmed to only use the green LED to allow for a refresh rate of 180 Hz. The emitted light from the projectors was long-pass filtered to restrict it to wavelengths above 550 nm, allowing for a maximal luminance of 250 ± 68 cd m^−2^ (mean ± SD). Stimuli were rendered using custom-written software in Python (v.2.7). Panda3D (v.1.8) was used to pre-distort the images to account for the curvature of the screen. Reported contrast values represent the Michelson contrast:

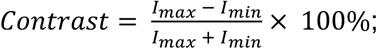

where *I*_max a_nd *I*_min r_epresent the maximal and minimal pixel intensity values (8-bit) of the stimulus.

### Histamine imaging

To measure changes in histamine concentrations in the extracellular space surrounding the dendrites of L1, L2, T4, and T5 neurons in the same fly, flies bearing a split-*GAL4* driver for L1 and L2 neurons were crossed to flies bearing a T4/T5-*GAL4* driver and the genetically encoded histamine sensor HA1m.^41,72^ L1 and L2 neurons were stimulated by displaying full-field 100% contrast ON–OFF flashes of three different durations (2, 4, and 10 s in total, first half ON, second half OFF), in pseudorandom order over 10 repetitions. T4 neurons were stimulated by displaying 50% contrast ON edges moving over the screen at a velocity of 50° s^−1^ in 8 different directions separated by 45° over 5 repetitions, also presented in pseudorandom order. T5 neurons were stimulated similarly, but using OFF edges. All three fields of view (lamina, M10, and Lo1) were imaged consecutively in each fly and ΔF/F signals were averaged over stimulus repetitions.

### Calcium imaging

Changes in intracellular calcium concentration in T4 and T5 neurons were measured using the genetically encoded calcium sensor *GCaMP8m*,^53^ driven by a *GAL4* line targeting a- and d-subtypes (T4/T5ad > *GCaMP8m*). To investigate the effect of silencing different genes on the visual response properties of T4 and T5 neurons, flies expressing T4/T5ad > *GCaMP8m* were crossed to flies bearing constructs for targeted RNAi. Two types of control flies were used. The first did not express any RNAi construct (control) and was generated by crossing flies of the T4/T5ad > *GCaMP8m* genotype to Canton S wild-type flies. The second expressed two different RNAi constructs for interference with *HisCl1* to provide a control that harbors at least as many RNAi transgenes as the experimental groups.^52^ Effects of knocking down *Rdl* and *ort* separately and in conjunction were tested using RNAi constructs targeting the respective transcripts.^50,51^

Responses to full-field flashes were imaged in M10 (T4ad) and Lo1 (T5ad). The stimulus consisted of 50% contrast ON–OFF flashes (2s ON, 2s OFF) and was repeated 5 times. ΔF/F values were averaged over repetitions for each ROI and over flies for each genotype. Responses were quantified as maximal ΔF/F during the 300 ms following each flash.

Fine-grained directional tuning curves of T4a neurons were obtained by recording in lobula plate layer 1, using a visual stimulation protocol consisting of ON edges (50% contrast) moving over the screen at 50° s^−1^ in 20 different directions separated by 18° in pseudorandom order over three repetitions. T5 neurons were recorded in the same field of view, but using OFF edges. ΔF/F signals of each ROI were averaged over stimulus repetitions. Because all recorded neurons receive visual input from different parts of the stimulus screen, signals from different ROIs could not be averaged based on the start time of the stimulus. Instead, the maximal ΔF/F was determined within the 5.18 s that each stimulus lasted, for each ROI and each stimulus direction separately. This peak time was taken as a point of reference to align edge responses across ROIs. To calculate the preferred direction and the directional tuning index (*L*_dir)_, the minimum ΔF/F value of each ROI was subtracted from the peak values, resulting in 20 vectors ***v***(*ϕ*) representing positive responses to each stimulus direction (*ϕ*). The preferred direction of each ROI was taken to be the stimulus direction closest to the direction of the vector sum of the 20 response vectors. *L*_dir w_as calculated by dividing the magnitude of the vector sum by the sum of all 20 individual response vector magnitudes:

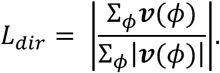

*L*_dir c_an vary between 0 and 1, representing the lowest possible directional tuning (the same response to stimuli in all directions) and the highest possible directional tuning (no response to stimuli in any direction but one), respectively. Averaging over the three ROIs of each fly and over the flies of each genotype was done based on the relative angular distance of each stimulus to the individually calculated preferred direction. Responses were normalized to the maximal ΔF/F of each ROI. Polar plots of maximal ΔF/F were normalized again to the maximum of each genotype, to make the shapes comparable between genotypes.

The tuning strength of T4a and T5a neurons to stimulus polarity was quantified from calcium traces in M10 and Lo1 in response to 50% contrast, 50° s^−1^ velocity ON and OFF edges moving in the respective preferred direction, repeated 5 times. Traces were averaged over repetitions and aligned to their peaks as above for directional tuning, but were not normalized. Response values represent the maximal ΔF/F during the stimulus. The polarity selectivity index (*PSI*) was quantified for each ROI based on its maximal response to ON and OFF stimuli:

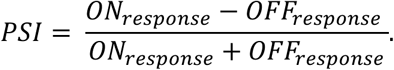

The *PSI* has a theoretical range between −1 and 1, with 1 representing a ROI that responds only to ON edges and −1 representing a ROI that responds only to OFF edges.

### Dendrite rendering

Using the fafbseg (v.3.2)^14–16^ and navis (v.1.12)^80^ packages in Python (v.3.11), the dendrite meshes of a T4c (cell ID 720575940612221669) and T5c (cell ID 720575940627480842) neuron were loaded from FAFB v783 and plotted using the matplotlib (v.3.10) backend, together with postsynapses that were colored in accordance to ultrastructure-based neurotransmitter predictions of the corresponding presynaptic neurons.^17^

### RNA sequencing analysis

Cell-type specific expression values of genes of interest were extracted from four published sequencing datasets and averaged over developmental stages and T4/T5 subtypes.^24–27^ For data from Özel et al.,^26^ adult data were plotted separately from developmental data. For data from Yoo et al.,^27^ the DGRP dataset was used. The plotted values represent log-transformed transcripts per 10,000 (TP10K) or unique molecular identifiers (UMI), except for data from Davis et al.,^25^ from which expression probability values were plotted.

### Biophysical model

Model simulations were performed using the Jaxley library in Python (v.3.14).^78^ The morphology of a T4 neuron (cell ID 720575940612221669, FAFB v783)^14–16^ was discretized using the *dλ* rule^85^ with *dλ* = 0.1. Consistent with input resistance measurements from T4 cells^20^ and standard values for invertebrate neurons,^86,87^ the compartmental model had a specific membrane resistance of 20 kΩ·cm^2^, an axial resistivity of 250 Ω·cm, and a specific membrane capacitance of 1 µF cm^−2^. In addition, the axonal compartments of the model contained a T4/T5 cell-specific voltage-gated sodium conductance.^56^ Simulations were performed with a time step of 25 µs.

Dendritic synapses were placed at their precise subcellular locations (Figure 1A) and grouped by neurotransmitter (glutamate, acetylcholine, and GABA). Each group was connected to one artificial single-compartment neuron whose activity mimicked the response properties of the respective glutamatergic, cholinergic, and GABAergic presynaptic neurons.^20^ Synaptic conductances (*ḡ*_s)_ were modelled as functions of presynaptic membrane voltage (*V*_m)_:

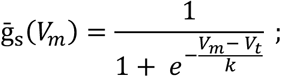

with a threshold voltage (*V*_t)_ of −57 mV and a slope (*k*) of 2 mV. Synaptic reversal potentials were set to −70 mV for glutamatergic synapses, −21 mV for cholinergic synapses, and −68 mV for GABAergic synapses, in accordance with empirical measurements.^20^ To simulate the voltage-dependent component of Rdl, all compartments with GABAergic synapses were also equipped with a voltage-gated chloride channel, whose current-voltage relationship and activation kinetics were fit to voltage-clamp recordings from Rdl-expressing S2R+ cells (Figures 1H and 1I).

Gaussian current pulses were injected into presynaptic neurons in distinct temporal sequences to simulate a bright bar moving at a velocity (*v*) of 25° s^−1^. For a given angle *φ* of visual motion, the delay between input signals was calculated as Δ*t* = *θ* ·cos*φ*/*v* assuming an inter-ommatidial angle *θ* of 5°. A delay of 200 ms was added to the glutamatergic and the GABAergic input arms, which were additionally low-pass filtered by increasing the membrane capacitance to capture the response dynamics of the presynaptic neurons.^20,55^ Directional tuning of the T4 neuron was assessed by simulating motion in 20 different directions separated by 18° and calculating the average membrane potential deflection (at the axon terminal) over a peak-centered 150-ms time window relative to the resting membrane potential. Three scenarios were compared: The complete model including both ligand- and voltage-gating of Rdl; a model without the voltage-gated component of Rdl; and a model completely devoid of Rdl. Results were normalized to the preferred-direction response of the respective models.

### Behavior

Female flies (1–5 days old) were cold-immobilized and attached to a pin with a drop of UV-curing dental glue (Sinfony Opaque Dentin, 3M) connecting the head and thorax. The wings were also glued to the pin. Each tethered fly was positioned on top of a 6 mm (diameter) air-suspended ball in a recording setup described before^57,88^ and was allowed to walk freely in all virtual directions by rotating the ball with its legs. The rotation of the ball was tracked, using a custom-designed system consisting of two optical computer mouse sensors, and digitized at 200 Hz. The air temperature around the fly was kept at 34 °C during the experiment to encourage flies to walk, using a custom-built air conditioning device including a Peltier heater (Quick-Cool, Cat#QC-127-1.4-6.0MS). Three displays (Samsung, Cat#2233RZ) with a maximal luminance of 131 cd m^−2^ were positioned vertically to form a U-shaped visual arena around the fly, spanning 270° in azimuth and 120° in elevation of the fly’s visual field. Visual stimuli were presented at 50% contrast and a refresh rate of 120 Hz. Custom-written software in Python (v.2.7) and Panda3D (v.1.8) was used to pre-distort the images to create a cylindrical panorama.

In open-loop experiments, the visual stimuli contained single ON or OFF edges, that traversed the visual arena in the right- or leftwards direction at 60° s^−1^. The first 15 of 50 trials were used to equilibrate temperature and were excluded from data analysis. Additional inclusion criteria were a forward walking speed of ≥ 0.15 cm s^−1^ (on a trial-by-trial basis) and a minimum of ten remaining trials per fly. To correct for rotational biases, the average azimuthal rotation in every full trial (including right- and leftward stimulation) was subtracted from each individual trace. The optomotor response strength was quantified as a fly’s rotational velocity in the direction of the visual stimulus, and was averaged over flies and genotypes. Average and peak responses to back-to-front and front-to-back stimulation were extracted from the time spans during which the edge traversed two 80° regions on the left and right side of the visual field, which were placed adjacent to the 20° frontal region.

Closed-loop experiments were carried out as described before.^20^ The azimuthal position of a 10°-wide dark vertical bar was controlled by the rotation of the ball in real time (updated every ∼9 ms), with the change in bar position indirectly proportional to the rotation about the z-axis. The bar appeared at a random azimuthal position at the start of each trial. During 180 trials of 20 s, the fly could control the bar position through its walking behavior. The first 40 trials were used to equilibrate temperature and were excluded from data analysis. Additional inclusion criteria were a forward walking speed of ≥ 0.30 cm s^−1^ (on a trial-by-trial basis), a minimum of 50 remaining trials per fly, and a rotational bias < 10° s^−1^. The probability density function of bar positions was calculated for each trial using a bin width of 5° before averaging over trials. Fixation in front was quantified by summing the probabilities of finding the bar in the 60° window in front of the fly and averaging these probabilities over trials.

### Quantification and statistical analysis

Statistical tests were performed in Prism (v.11.1, GraphPad). Details of all statistical tests (name, measured variable, experimental groups/comparisons, test statistic, and *P*-value) are compiled in Tables S1 and S2. *P*-values < 0.05 were considered significant. Exact *n*-values are reported in the figures; the corresponding figure legends describe what *n* represents. In cases where no statistical tests were performed, the number of biological replicates is stated in the figure legend. Definitions of center and dispersion are stated in the results and figure legends, wherever applicable. Data were assessed for normality using the Shapiro–Wilk test and for equality of variances using the Brown–Forsythe test. Two groups of normally distributed data were compared using two-tailed Student’s *t*-tests. If one of the groups did not pass the normality test or had a low (*n* < 5) sample size, the nonparametric two-tailed Mann–Whitney *U*-test was used instead. Differences between the means of multiple independent groups of data that met the assumptions of normality and equality of variances were compared using one-way analysis of variance (ANOVA). Where the assumption of normality was violated, group medians were compared using Kruskal–Wallis test followed by Dunn’s multiple-comparisons test. Multiple-comparisons tests were only performed after a significant Kruskal–Wallis test and included only predefined comparisons of interest. Reported *P*-values were corrected for multiple comparisons. For multiple comparisons with two parental controls, the highest of two *P*-values is reported in the figure legend.

---

**Figure S1.**
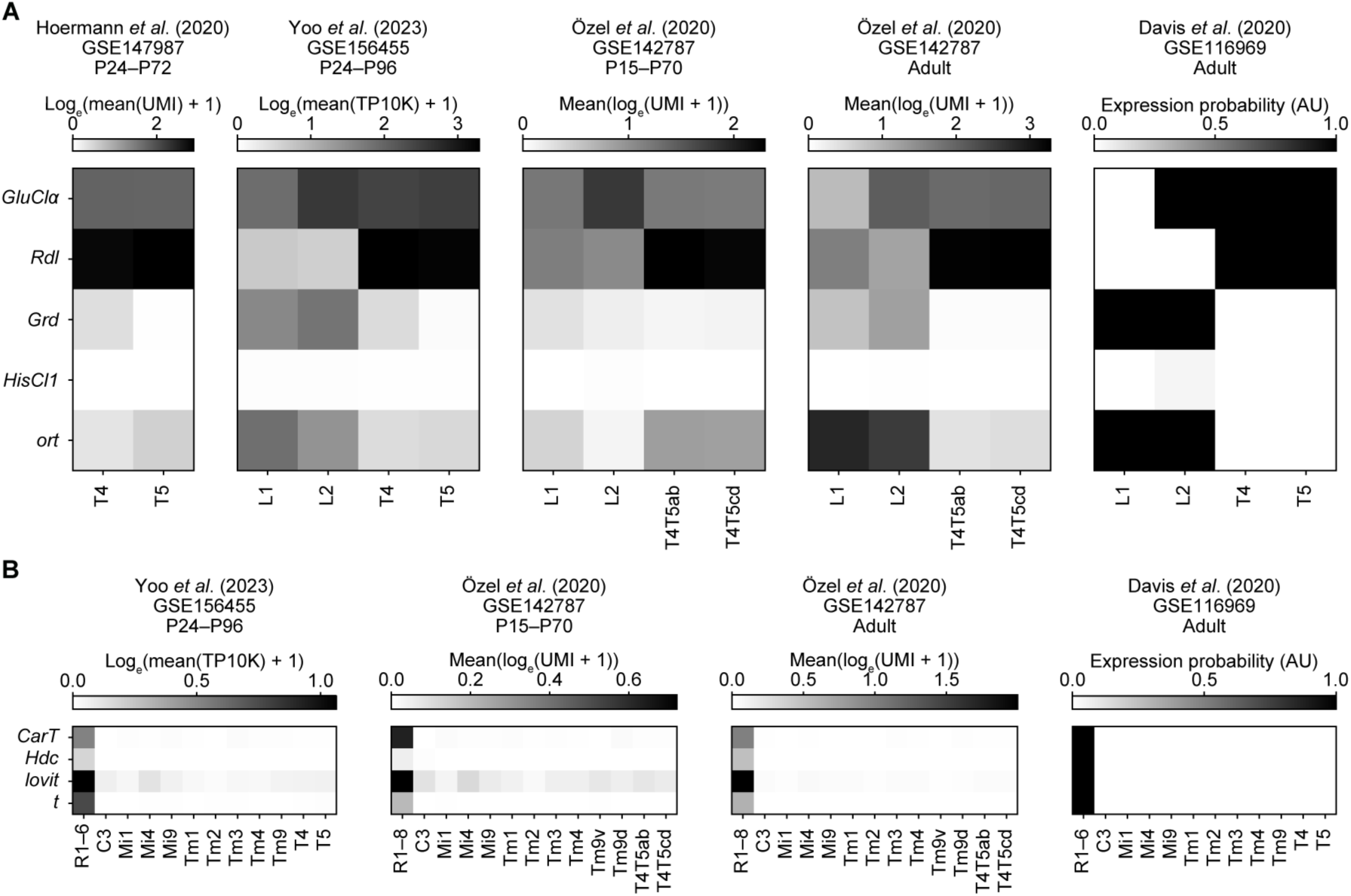
RNA sequencing datasets of cys-loop ligand-gated receptors and histamine-related genes. (A) Heatmaps of RNA expression values of cys-loop ligand-gated chloride channels in developing and adult neurons, from RNA sequencing data (GEO: GSE147987,^24^ GSE156455,^27^ GSE142787,^26^ and GSE116969).^25^ Developmental data are averaged over the time points specified. Data corresponding to (GSE156455)^27^ are replicated from Figure 1C for reference. (B) Heatmaps of RNA expression values of genes involved in histamine synthesis, transport, and recycling, in developing and adult T4 and T5 neurons and their direct inputs, as well as in the known histaminergic photoreceptor (R) neurons, with similar quantification as (A).^25–27^ See also Figure 1 and STAR Methods.

**Figure S2.**
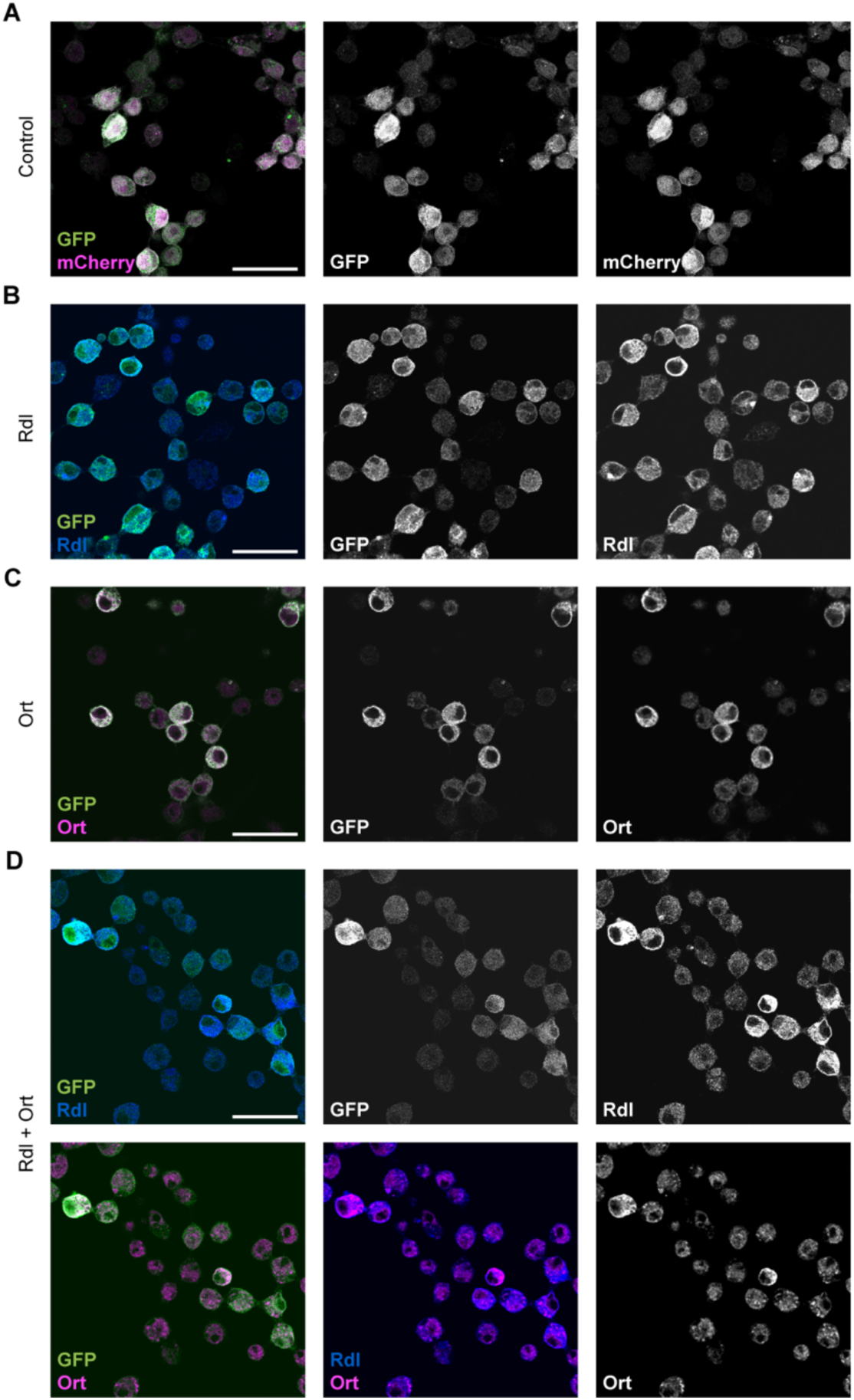
S2R+ immunocytochemistry. Micrographs of cell cultures transfected with plasmid to express (A) *EGFP* and *mCherry* (control), (B) *Rdl* and *EGFP* (Rdl), (C) *ort::EGFP* (Ort), or (D) *Rdl*, *ort* and *EGFP* (Rdl + Ort), stained with antibodies against GFP (green), Ort (magenta), and Rdl (blue). Merged channels show two colors together and single channels are displayed in grayscale. Micrographs are representative of three separately transfected coverslips. Scale bars, 20 µm. See also Figure 1 and STAR Methods.

**Figure S3.**
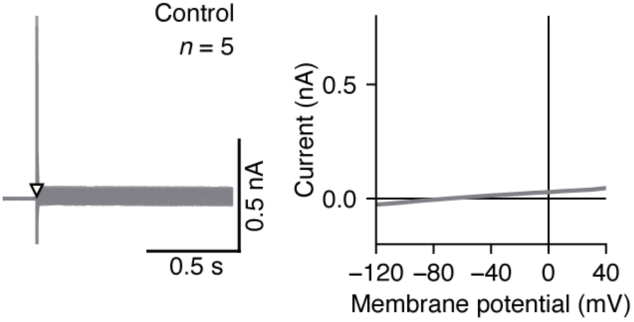
Control cells do not exhibit voltage-gated currents. Average currents elicited by voltage steps from a clamp potential of −70 to potentials between −120 and +40 mV (left, empty triangle) in cells expressing the control plasmid, and the corresponding current-voltage relationship (right, mean ± SEM). Data are analyzed from the same recordings as displayed in Figure 2H, with 50 mM GABA applications 1.05 s after the voltage step (not shown here). *n* value indicates the number of cells. See also Figure 1 and STAR Methods.

**Figure S4.**
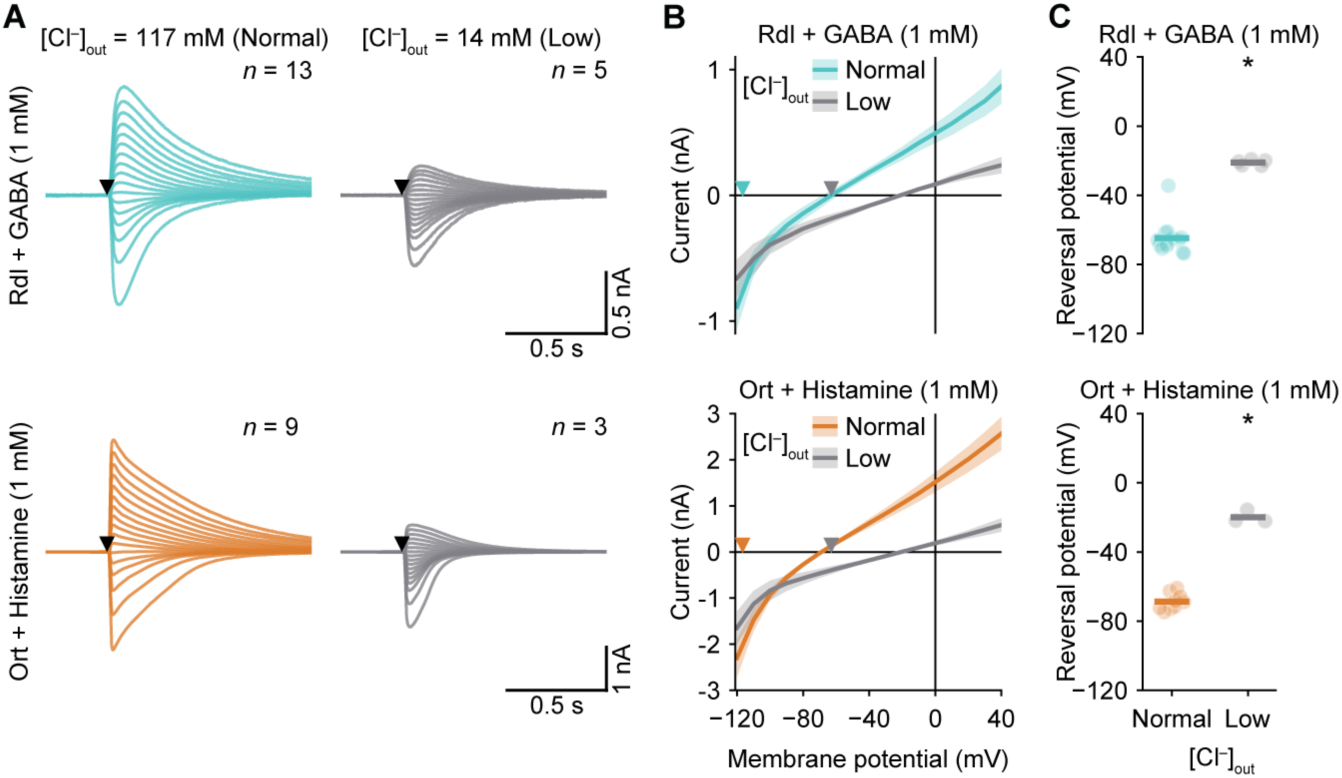
**Extracellular chloride concentration affects the reversal potential**(A) Average currents elicited by applications (filled triangles) of 1 mM GABA in *Rdl*-expressing cells (top) or 1 mM histamine in *ort*-expressing cells (bottom) at 17 clamp potentials ranging from −120 to +40 mV, with an extracellular chloride concentration of 117 mM (normal, left) or 14 mM (low, right). Data corresponding to 117 mM chloride are also used in Figure 1H and 1I. (B) Current-voltage relationships (mean ± SEM) of the data in (A). Filled triangles indicate theoretical reversal potentials of chloride ions corresponding to the normal (cyan and orange) and low (gray) extracellular concentration. (C) Average (horizontal lines) reversal potentials of all conditions in (A). Light circles represent individual cells. Significantly different reversal potentials from the normal chloride concentration, determined using two-tailed Mann–Whitney U-tests, are indicated; \**P* = 2.3 × 10^−4^ (Rdl + GABA), \**P* = 9.1 × 10^−3^ (Ort + Histamine). *n* values indicate the number of cells. See also Figure 1, Table S2, and STAR Methods.

**Figure S5.**
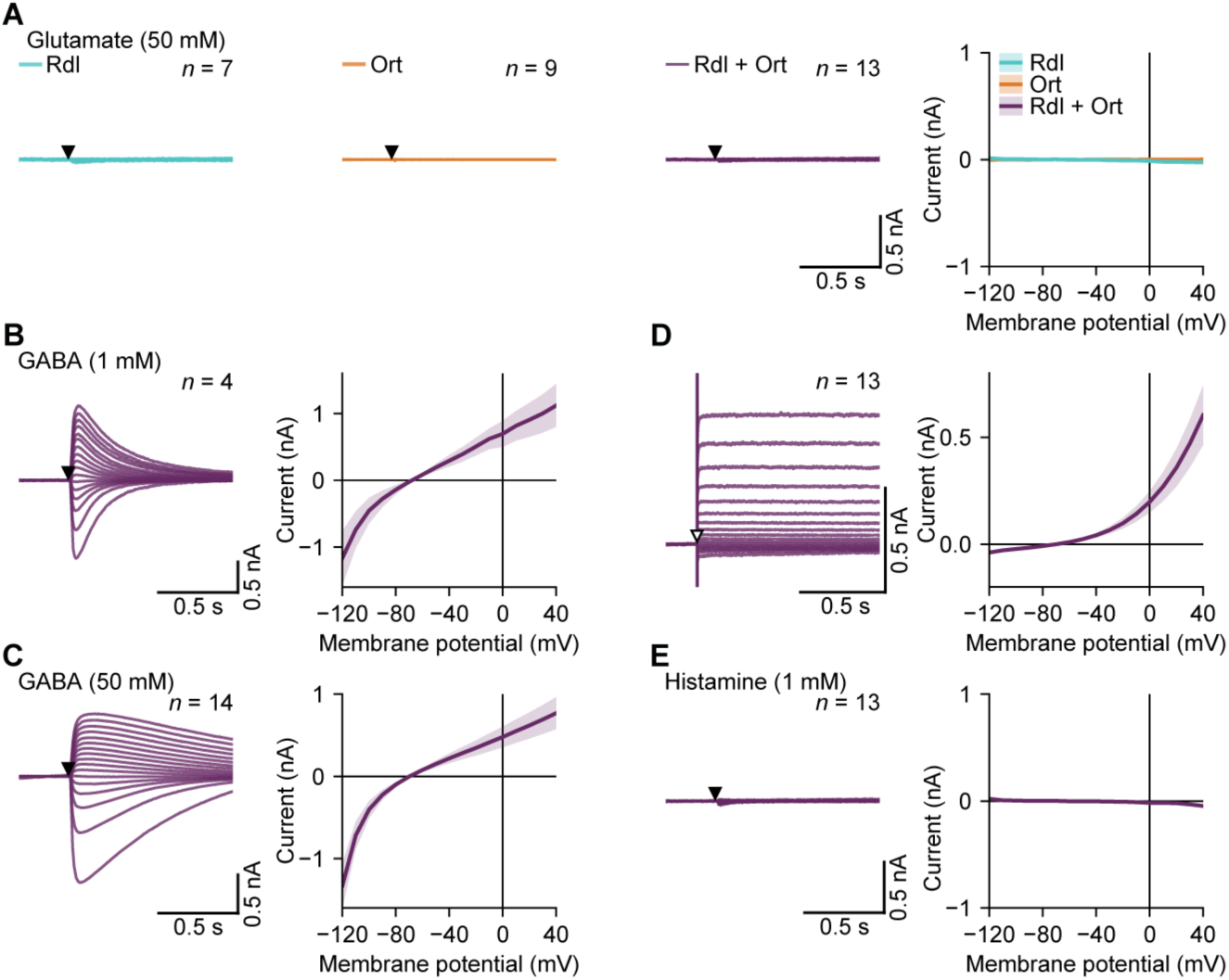
Glutamate does not elicit currents and cells transfected with *Rdl* and *ort* respond like *Rdl*-transfected cells. (A) Average currents (left) before and after applications (filled triangles) of 50 mM glutamate in cells expressing *Rdl* (cyan), *ort* (orange), or *Rdl* + *ort* (purple) at 17 clamp potentials ranging from −120 to +40 mV, and corresponding current-voltage relationships (mean ± SEM, right). (B–E) Average currents elicited by applications (filled triangles) of 1 mM GABA (B), 50 mM GABA (C), 1 mM histamine (E), or by voltage steps (D, empty triangles) in cells expressing *Rdl* + *ort* at 17 clamp potentials ranging from −120 to +40 mV (left), and the corresponding current-voltage relationships (right, mean ± SEM). *n* values indicate the number of cells. See also Figure 2 and STAR Methods.

**Figure S6.**
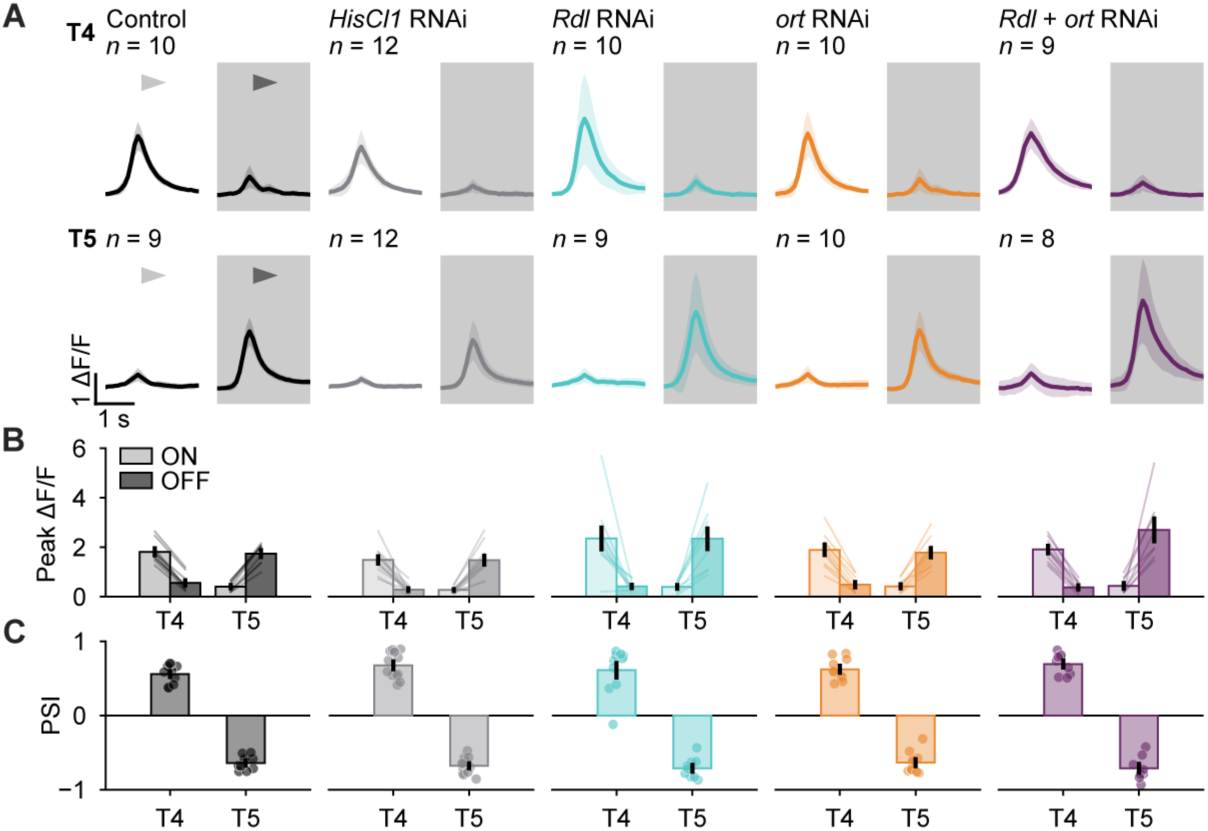
T4 and T5 neurons retain their ON/OFF selectivity when expressing RNAi constructs. (A) Calcium responses of T4ad (top) and T5ad (bottom) neurons in response to ON (white background) and OFF (gray background) edges of 50% contrast moving in the preferred direction of subtype a (arrowheads) with 50° s^−1^. Control neurons (black) expressed only *GCaMP8m*, the others additionally RNAi constructs to knock down *HisCl1* (gray), *Rdl* (cyan), *ort* (orange) or *Rdl* + *ort* (purple). Data are peak-aligned mean ± SD. (B) Maximal responses of data in (A). Bars are mean ± SEM. Light bars indicate responses to ON edges and dark bars to OFF edges. Lines connect responses of individual flies. (C) Polarity selectivity indices (PSI) of data in (B). No significant differences in PSI were found between the genotypes (T4: Kruskal–Wallis test, *P* = 4.1 × 10^−1^; T5: one-way ANOVA, *P* = 5.9 × 10^−1^). Bars represent mean ± SEM. Circles represent individual flies. *n* values indicate the number of flies. See also Figure 3, Tables S2 and S3, and STAR Methods.

**Figure S7.**
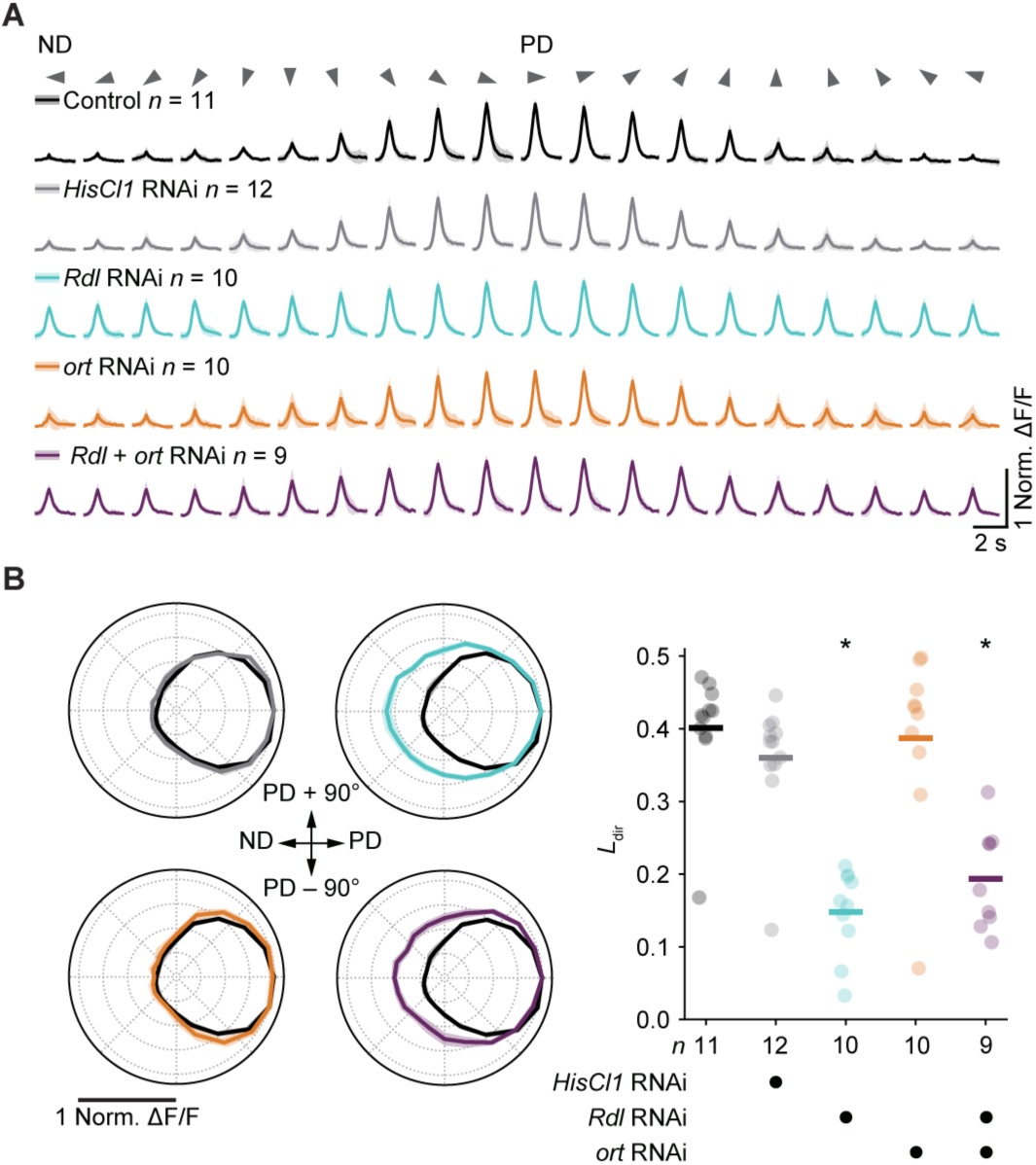
**T5 direction selectivity**(A) Normalized T5a calcium responses (mean ± SD) to OFF-edges moving at 50° s^−1^ in 20 different directions (arrowheads). Control neurons (black) expressed only *GCaMP8m*, the others additionally RNAi constructs to knock down *HisCl1* (gray), *Rdl* (cyan), *ort* (orange) or *Rdl* + *ort* (purple). (B) Data from (A) represented as polar plots (mean ± SEM) of the maximal responses (left) and corresponding *L*dir values (right). No-RNAi control data (black) are repeated in each polar plot for comparison. Significant differences in *L*dir compared to the no-RNAi control condition, determined using Dunn’s multiple comparisons test following Kruskal–Wallis test, are indicated; \**P* = 1.1 × 10^−4^ (*Rdl* RNAi), \**P* = 1.4 × 10^−3^ (*Rdl* + *ort* RNAi). PD, preferred direction; ND, null direction. *n* values indicate the number of flies. See also Figure 3, Tables S2 and S3, and STAR Methods.

**Figure S8.**
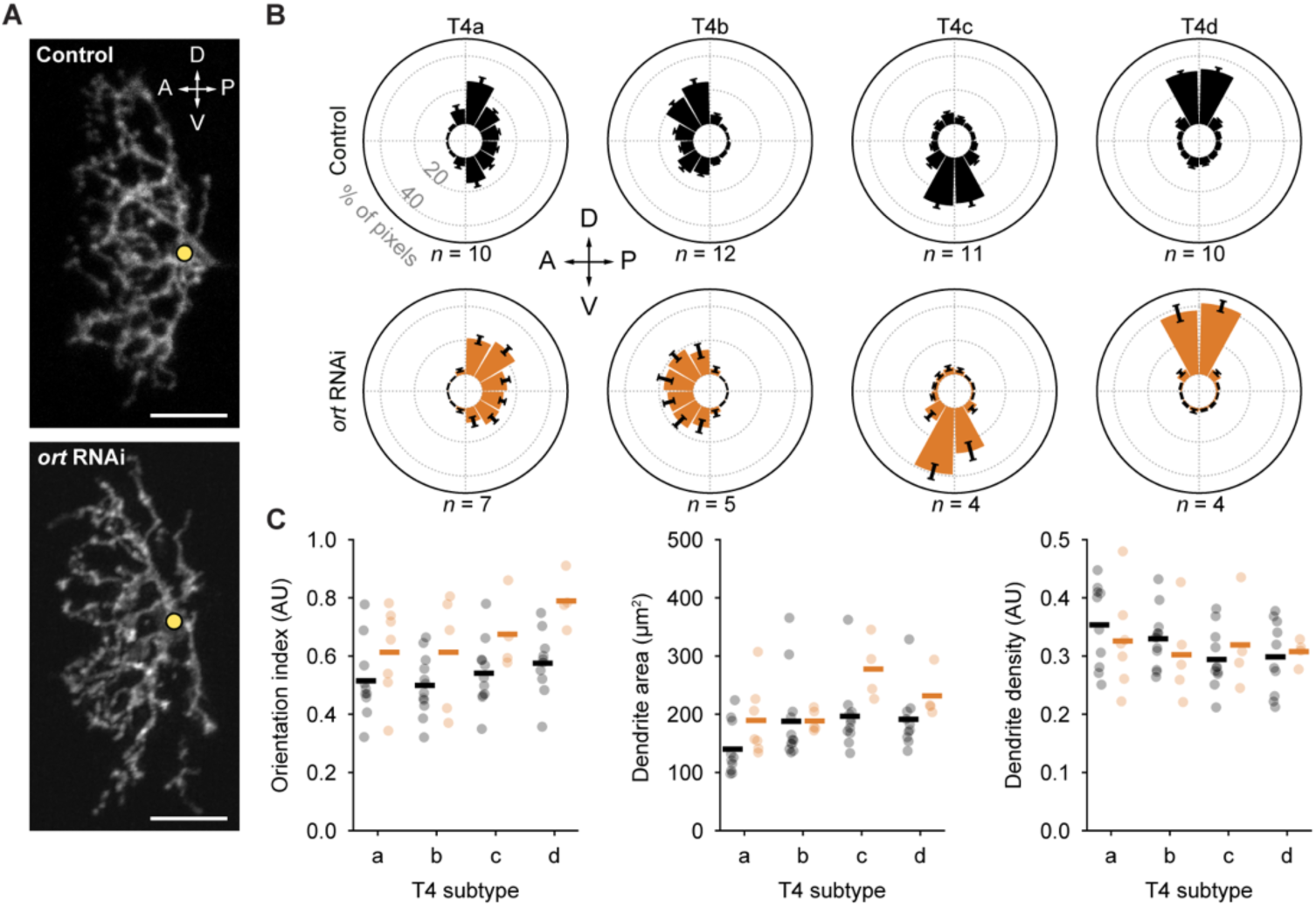
Dendrite morphologies of control and Ort-deficient T4 neurons. (A) Representative maximum intensity z-projections of dendrites from control (top) and *ort* RNAi-expressing (bottom) T4b neurons. The yellow circles indicate the dendrite origins. Scale bars, 5 µm. (B) Dendrite orientation histograms for all T4 subtypes of the control (black) and *ort* RNAi (orange) conditions. Bars represent the mean ± SEM percentage of dendrite pixels that occupy each 30° angular region from the dendrite origin. (C) Orientation index (left), dendrite area (middle), and dendrite density (right) of all dendrites that were also analysed in (B). Dunn’s multiple comparisons test following Kruskal–Wallis test (orientation index and dendrite area) or Kruskal–Wallis test (dendrite density) did not find significant differences between control and Ort-deficient dendrites for any of the subtypes (see Table S2). Horizontal lines represent averages and circles indicate individual neurons. A, anterior; P, posterior; D, dorsal; V, ventral; AU, arbitrary units. *n* values indicate the number of cells. See also Figure 3, Tables S2 and S3, and STAR Methods.

## Notes

### Competing Interest Statement

The authors have declared no competing interest.

