## Supplementary Tables for "Non-canonical inhibition in motion vision"

### Table S1. Statistical analyses for Figures 2, 3, and 5

| Fig. | Statistical test | Measured variable (unit) | Experimental groups/comparisons | Test statistic | *P* |
|---|---|---|---|---|---|
| 2L | Shapiro–Wilk test | Exponential decay (τ) | *Rdl* | $W = 0.8269$ | 0.5806 |
| | Shapiro–Wilk test | Exponential decay (τ) | *ort* | $W = 0.9399$ | $3.068\times10^{-2}$ |
| | Two-tailed Mann–Whitney *U*-test | Exponential decay (τ) | *Rdl* **vs.** *ort* | $U = 0$ | $2.165\times10^{-5}$ |
| 3C | Shapiro–Wilk test | *Rdl* T4 norm. intensity (AU) | *HisCl1* RNAi | $W = 0.9040$ | 0.3137 |
| | Shapiro–Wilk test | *Rdl* T4 norm. intensity (AU) | *Rdl* RNAi | $W = 0.9207$ | 0.5101 |
| | Two-tailed unpaired Student's *t*-test | *Rdl* T4 norm. intensity (AU) | *HisCl1* RNAi **vs.** *Rdl* RNAi | $t_{12} = 3.800$ | $2.531\times10^{-3}$ |
| | Shapiro–Wilk test | *Rdl* T5 norm. intensity (AU) | *HisCl1* RNAi | $W = 0.9697$ | 0.8953 |
| | Shapiro–Wilk test | *Rdl* T5 norm. intensity (AU) | *Rdl* RNAi | $W = 0.9627$ | 0.8401 |
| | Two-tailed unpaired Student's *t*-test | *Rdl* T5 norm. intensity (AU) | *HisCl1* RNAi **vs.** *Rdl* RNAi | $t_{12} = 5.621$ | $1.124\times10^{-4}$ |
| | Shapiro–Wilk test | *ort* T4 norm. intensity (AU) | *HisCl1* RNAi | $W = 0.9681$ | 0.8632 |
| | Shapiro–Wilk test | *ort* T4 norm. intensity (AU) | *ort* RNAi | $W = 0.8932$ | 0.3351 |
| | Two-tailed unpaired Student's *t*-test | *ort* T4 norm. intensity (AU) | *HisCl1* RNAi **vs.** *ort* RNAi | $t_9 = 8.712$ | $1.113\times10^{-5}$ |
| | Shapiro–Wilk test | *ort* T5 norm. intensity (AU) | *HisCl1* RNAi | $W = 0.8532$ | 0.2049 |
| | Shapiro–Wilk test | *ort* T5 norm. intensity (AU) | *ort* RNAi | $W = 0.9189$ | 0.4976 |
| | Two-tailed unpaired Student's *t*-test | *ort* T5 norm. intensity (AU) | *HisCl1* RNAi **vs.** *ort* RNAi | $t_9 = 8.660$ | $1.168\times10^{-5}$ |
| 3E | Shapiro–Wilk test | T4 max flash response (ΔF/F) | Control | $W = 0.8553$ | $6.712\times10^{-2}$ |
| | Shapiro–Wilk test | T4 max flash response (ΔF/F) | *HisCl1* RNAi | $W = 0.8051$ | $1.099\times10^{-2}$ |
| | Shapiro–Wilk test | T4 max flash response (ΔF/F) | *Rdl* RNAi | $W = 0.8292$ | $5.722\times10^{-2}$ |
| | Shapiro–Wilk test | T4 max flash response (ΔF/F) | *ort* RNAi | $W = 0.7116$ | $1.216\times10^{-3}$ |
| | Shapiro–Wilk test | T4 max flash response (ΔF/F) | *Rdl* + *ort* RNAi | $W = 0.9285$ | 0.4674 |
| | Kruskal–Wallis test | T4 max flash response (ΔF/F) | All groups | $H = 30.04$ | $4.802\times10^{-6}$ |
| | Dunn's multiple comparisons test | T4 max flash response (ΔF/F) | Control **vs.** *HisCl1* RNAi | $Z = 0.6717$ | > 0.9999 |
| | | | Control **vs.** *Rdl* RNAi | $Z = 3.264$ | $4.395\times10^{-3}$ |
| | | | Control **vs.** *ort* RNAi | $Z = 0.3514$ | > 0.9999 |
| | | | Control **vs.** *Rdl* + *ort* RNAi | $Z = 3.458$ | $2.176\times10^{-3}$ |
| | Shapiro–Wilk test | T5 max flash response (ΔF/F) | Control | $W = 0.9846$ | 0.9839 |
| | Shapiro–Wilk test | T5 max flash response (ΔF/F) | *HisCl1* RNAi | $W = 0.9760$ | 0.9401 |
| | Shapiro–Wilk test | T5 max flash response (ΔF/F) | *Rdl* RNAi | $W = 0.8063$ | $3.338\times10^{-2}$ |
| | Shapiro–Wilk test | T5 max flash response (ΔF/F) | *ort* RNAi | $W = 0.8464$ | $6.806\times10^{-2}$ |
| | Shapiro–Wilk test | T5 max flash response (ΔF/F) | *Rdl* + *ort* RNAi | $W = 0.9220$ | 0.4464 |
| | Kruskal–Wallis test | T5 max flash response (ΔF/F) | All groups | $H = 13.92$ | $7.540\times10^{-3}$ |
| | Dunn's multiple comparisons test | T5 max flash response (ΔF/F) | Control **vs.** *HisCl1* RNAi | $Z = 0.7546$ | > 0.9999 |
| | | | Control **vs.** *Rdl* RNAi | $Z = 0.7639$ | > 0.9999 |
| | | | Control **vs.** *ort* RNAi | $Z = 0.5204$ | > 0.9999 |
| | | | Control **vs.** *Rdl* + *ort* RNAi | $Z = 2.527$ | $4.606\times10^{-2}$ |
| 3G | Shapiro–Wilk test | $L_{dir}$ (AU) | Control | $W = 0.8914$ | 0.1450 |
| | Shapiro–Wilk test | $L_{dir}$ (AU) | *HisCl1* RNAi | $W = 0.8120$ | $1.285\times10^{-2}$ |
| | Shapiro–Wilk test | $L_{dir}$ (AU) | *Rdl* RNAi | $W = 0.9037$ | 0.2405 |
| | Shapiro–Wilk test | $L_{dir}$ (AU) | *ort* RNAi | $W = 0.8396$ | $3.123\times10^{-2}$ |
| | Shapiro–Wilk test | $L_{dir}$ (AU) | *Rdl* + *ort* RNAi | $W = 0.9379$ | 0.5603 |
| | Kruskal–Wallis test | $L_{dir}$ (AU) | All groups | $H = 33.59$ | $9.063\times10^{-7}$ |
| | Dunn's multiple comparisons test | $L_{dir}$ (AU) | Control **vs.** *HisCl1* RNAi | $Z = 1.511$ | 0.5229 |
| | | | Control **vs.** *Rdl* RNAi | $Z = 4.447$ | $3.478\times10^{-5}$ |
| | | | Control **vs.** *ort* RNAi | $Z = 1.242$ | 0.8562 |
| | | | Control **vs.** *Rdl* + *ort* RNAi | $Z = 4.533$ | $2.327\times10^{-5}$ |
| 5A | Shapiro–Wilk test | Peak rot. vel. FB − BF (° $s^{-1}$) | T4/T5 | $W = 0.9297$ | $6.785\times10^{-2}$ |
| | Shapiro–Wilk test | Peak rot. vel. FB − BF (° $s^{-1}$) | *Rdl* RNAi | $W = 0.9463$ | 0.2452 |
| | Shapiro–Wilk test | Peak rot. vel. FB − BF (° $s^{-1}$) | T4/T5 > *Rdl* RNAi | $W = 0.8891$ | $8.996\times10^{-3}$ |
| | Shapiro–Wilk test | Peak rot. vel. FB − BF (° $s^{-1}$) | *ort* RNAi | $W = 0.9326$ | 0.1244 |
| | Shapiro–Wilk test | Peak rot. vel. FB − BF (° $s^{-1}$) | T4/T5 > *ort* RNAi | $W = 0.9511$ | 0.3566 |
| | Shapiro–Wilk test | Peak rot. vel. FB − BF (° $s^{-1}$) | *Rdl* + *ort* RNAi | $W = 0.8950$ | $2.804\times10^{-2}$ |
| | Shapiro–Wilk test | Peak rot. vel. FB − BF (° $s^{-1}$) | T4/T5 > *Rdl* + *ort* RNAi | $W = 0.9571$ | 0.4326 |
| | Kruskal–Wallis test | Peak rot. vel. FB − BF (° $s^{-1}$) | All groups | $H = 56.99$ | $1.833\times10^{-10}$ |
| | Dunn's multiple comparisons test | Peak rot. vel. FB − BF (° $s^{-1}$) | T4/T5 **vs.** T4/T5 > *Rdl* RNAi | $Z = 5.244$ | $9.418\times10^{-7}$ |
| | | | *Rdl* RNAi **vs.** T4/T5 > *Rdl* RNAi | $Z = 4.249$ | $1.291\times10^{-4}$ |
| | | | T4/T5 **vs.** T4/T5 > *ort* RNAi | $Z = 2.474$ | $8.010\times10^{-2}$ |
| | | | *ort* RNAi **vs.** T4/T5 > *ort* RNAi | $Z = 1.567$ | 0.7025 |
| | | | T4/T5 **vs.** T4/T5 > *Rdl* + *ort* RNAi | $Z = 5.658$ | $9.206\times10^{-8}$ |
| | | | *Rdl* + *ort* RNAi **vs.** T4/T5 > *Rdl* + *ort* RNAi | $Z = 3.697$ | $1.307\times10^{-3}$ |
| 5B | Shapiro–Wilk test | Fixation in front (%) | T4/T5 | $W = 0.9822$ | 0.9772 |
| | Shapiro–Wilk test | Fixation in front (%) | *Rdl* RNAi | $W = 0.9158$ | 0.2201 |
| | Shapiro–Wilk test | Fixation in front (%) | T4/T5 > *Rdl* RNAi | $W = 0.9377$ | 0.5577 |
| | Shapiro–Wilk test | Fixation in front (%) | *ort* RNAi | $W = 0.9494$ | 0.6831 |
| | Shapiro–Wilk test | Fixation in front (%) | T4/T5 > *ort* RNAi | $W = 0.9790$ | 0.9475 |
| | Shapiro–Wilk test | Fixation in front (%) | *Rdl* + *ort* RNAi | $W = 0.8339$ | $1.771\times10^{-2}$ |
| | Shapiro–Wilk test | Fixation in front (%) | T4/T5 > *Rdl* + *ort* RNAi | $W = 0.9828$ | 0.9753 |
| | Kruskal–Wallis test | Fixation in front (%) | All groups | $H = 38.42$ | $9.303\times10^{-7}$ |
| | Dunn's multiple comparisons test | Fixation in front (%) | T4/T5 **vs.** T4/T5 > *Rdl* RNAi | $Z = 3.682$ | $1.390\times10^{-3}$ |
| | | | *Rdl* RNAi **vs.** T4/T5 > *Rdl* RNAi | $Z = 3.779$ | $9.458\times10^{-4}$ |
| | | | T4/T5 **vs.** T4/T5 > *ort* RNAi | $Z = 1.804$ | 0.4276 |
| | | | *ort* RNAi **vs.** T4/T5 > *ort* RNAi | $Z = 2.678$ | $4.444\times10^{-2}$ |
| | | | T4/T5 **vs.** T4/T5 > *Rdl* + *ort* RNAi | $Z = 3.723$ | $1.179\times10^{-3}$ |
| | | | *Rdl* + *ort* RNAi **vs.** T4/T5 > *Rdl* + *ort* RNAi | $Z = 3.166$ | $9.280\times10^{-3}$ |

#### Table S2. Statistical analyses for Figures S4–S8

| Fig. | Statistical test | Measured variable (unit) | Experimental groups/comparisons | Test statistic | *P* |
|---|---|---|---|---|---|
| **S4** | Shapiro–Wilk test | *Rdl* reversal potential (mV) | normal $[Cl^-]_{out}$ | *W* = 0.6940 | $4.836\times10^{-4}$ |
| | Shapiro–Wilk test | *Rdl* reversal potential (mV) | low $[Cl^-]_{out}$ | *W* = 0.8846 | 0.3305 |
| | Two-tailed Mann–Whitney *U*-test | *Rdl* reversal potential (mV) | normal $[Cl^-]_{out}$ **vs.** low $[Cl^-]_{out}$ | *U* = 0 | $2.334\times10^{-4}$ |
| | Two-tailed Mann–Whitney *U*-test | *ort* reversal potential (mV) | normal $[Cl^-]_{out}$ **vs.** low $[Cl^-]_{out}$ | *U* = 0 | $9.091\times10^{-3}$ |
| **S6** | Shapiro–Wilk test | T4 PSI (AU) | Control | *W* = 0.8685 | $9.605\times10^{-2}$ |
| | Shapiro–Wilk test | T4 PSI (AU) | *HisCl1* RNAi | *W* = 0.8869 | 0.1075 |
| | Shapiro–Wilk test | T4 PSI (AU) | *Rdl* RNAi | *W* = 0.7843 | $9.341\times10^{-3}$ |
| | Shapiro–Wilk test | T4 PSI (AU) | *ort* RNAi | *W* = 0.9272 | 0.4206 |
| | Shapiro–Wilk test | T4 PSI (AU) | *Rdl* + *ort* RNAi | *W* = 0.9037 | 0.2746 |
| | Kruskal–Wallis test | T4 PSI (AU) | All groups | *H* = 3.990 | 0.4074 |
| | Shapiro–Wilk test | T5 PSI (AU) | Control | *W* = 0.9040 | 0.2758 |
| | Shapiro–Wilk test | T5 PSI (AU) | *HisCl1* RNAi | *W* = 0.9800 | 0.9838 |
| | Shapiro–Wilk test | T5 PSI (AU) | *Rdl* RNAi | *W* = 0.9250 | 0.4351 |
| | Shapiro–Wilk test | T5 PSI (AU) | *ort* RNAi | *W* = 0.8627 | $8.202\times10^{-2}$ |
| | Shapiro–Wilk test | T5 PSI (AU) | *Rdl* + *ort* RNAi | *W* = 0.9517 | 0.7282 |
| | Brown–Forsythe test | T5 PSI (AU) | All groups | $F_{4, 43}$ = 0.4506 | 0.7713 |
| | One-way ANOVA | T5 PSI (AU) | All groups | $F_{4, 43}$ = 0.7092 | 0.5901 |
| **S7** | Shapiro–Wilk test | $L_{dir}$ (AU) | Control | *W* = 0.6687 | $1.862\times10^{-4}$ |
| | Shapiro–Wilk test | $L_{dir}$ (AU) | *HisCl1* RNAi | *W* = 0.7065 | $9.740\times10^{-4}$ |
| | Shapiro–Wilk test | $L_{dir}$ (AU) | *Rdl* RNAi | *W* = 0.1787 | 0.1787 |
| | Shapiro–Wilk test | $L_{dir}$ (AU) | *ort* RNAi | *W* = 0.7757 | $7.337\times10^{-3}$ |
| | Shapiro–Wilk test | $L_{dir}$ (AU) | *Rdl* + *ort* RNAi | *W* = 0.9211 | 0.4015 |
| | Kruskal–Wallis test | $L_{dir}$ (AU) | All groups | *H* = 29.08 | $7.537\times10^{-6}$ |
| | Dunn's multiple comparisons test | $L_{dir}$ (AU) | Control **vs.** *HisCl1* RNAi | *Z* = 1.388 | 0.6606 |
| | | | Control **vs.** *Rdl* RNAi | *Z* = 4.193 | $1.102\times10^{-4}$ |
| | | | Control **vs.** *ort* RNAi | *Z* = 0.2210 | > 0.9999 |
| | | | Control **vs.** *Rdl* + *ort* RNAi | *Z* = 3.577 | $1.391\times10^{-3}$ |
| **S8** | Kruskal–Wallis test | Dendrite orientation (AU) | All groups | *H* = 16.64 | $1.987\times10^{-2}$ |
| | Dunn’s multiple comparisons test | Dendrite orientation (AU) | Control T4a **vs.** *ort* RNAi T4a | *Z* = 1.556 | 0.4787 |
| | | | Control T4b **vs.** *ort* RNAi T4b | *Z* = 1.471 | 0.5655 |
| | | | Control T4c **vs.** *ort* RNAi T4c | *Z* = 1.650 | 0.3958 |
| | | | Control T4d **vs.** *ort* RNAi T4d | *Z* = 2.061 | 0.1572 |
| | Kruskal–Wallis test | Dendrite area ($\mu m^2$) | All groups | *H* = 20.09 | $5.3860\times10^{-3}$ |
| | Dunn’s multiple comparisons test | Dendrite area ($\mu m^2$) | Control T4a **vs.** *ort* RNAi T4a | *Z* = 1.817 | 0.2768 |
| | | | Control T4b **vs.** *ort* RNAi T4b | *Z* = 0.8780 | > 0.9999 |
| | | | Control T4c **vs.** *ort* RNAi T4c | *Z* = 2.134 | 0.1313 |
| | | | Control T4d **vs.** *ort* RNAi T4d | *Z* = 1.485 | 0.5506 |
| | Kruskal–Wallis test | Dendrite density (AU) | All groups | *H* = 5.413 | 0.6097 |

#### Table S3. Fly genotypes

| Figure | Name | Genotype |
|---|---|---|
| **1D** | T4/T5 > *ort::HA*, *myr::GFP* (T4 analysis) | *P{hs-FLPG5.PEST}attP3/$w^-$; P{10XUAS-IVS-myr::GFP}attP40/CyO; P{UAS-ort.HA.spGFP11}/P{tubP-(FRT.GAL80)}VK00027, P{R42F06-GAL4}attp2* |
| | T4/T5 > *ort::HA*, *CD4::tdGFP* (T5 analysis) | *P{hs-FLPG5.PEST}attP3/$w^-$; P{R42F06-GAL4}su(Hw)attp5/P{UAS-CD4-tdGFP}8M2; P{UAS-ort.HA.spGFP11}/P{tubP-(FRT.GAL80)}VK00027, P{R42F06-GAL4}attp2* |
| **1E, 1F** | T4/T5 > *ort::HA, Rdl::GFP* | *$w^-/w^-$; P{20XUAS-Rdl.GFP}su(Hw)attP5/P{R42F06-GAL4}su(Hw)attP5; P{UAS-ort.HA.spGFP11}/+* |
| **2B** | L1/L2/T4/T5 > *HA1m* | *$w^+/w^-$; P{R42F06-GAL4}su(Hw)attP5/ P{R48A08-p65.AD}attP40; P{10xUAS-HA1m}VK00005/ P{R29G11-GAL4.DBD}attP2* |
| **2D** | T4/T5 > *Rdl::GFP;* Mi4 > *$Brp^{short}$::mCherry* | *$w^-/w^-$; P{8xLexAop2-$Brp^{short}$::mCherry}28E7/P{20XUAS-Rdl.GFP}su(Hw)attP5; P{R48A07-LexA}su(Hw)attP1/P{R42F06-GAL4}attp2* |
| **2E** | T4/T5 > *ort::HA;* Mi4 > *$Brp^{short}$::mCherry* | *$w^-/w^-$; P{R42F06-GAL4}su(Hw)attP5/P{8xLexAop2-$Brp^{short}$::mCherry}28E7; P{UAS-ort.HA.spGFP11}/P{R48A07-LexA}su(Hw)attP1* |
| **2F** | T4/T5 > *Rdl::GFP;* Tm3 > *$Brp^{short}$::mCherry* | *$w^-/w^-$; P{8xLexAop2-$Brp^{short}$::mCherry}28E7/P{20XUAS-Rdl.GFP}su(Hw)attP5; P{GMR13E12-lexA}su(Hw)attP1/P{R42F06-GAL4}attp2* |
| **2G** | T4/T5 > *ort::HA;* Tm3 > *$Brp^{short}$::mCherry* | *$w^-/w^-$; P{8xLexAop2-$Brp^{short}$::mCherry}28E7/P{R42F06-GAL4}su(Hw)attP5; P{GMR13E12-lexA}su(Hw)attP1/P{UAS-ort.HA.spGFP11}* |
| **3A, 3C** | *HisCl1* RNAi | *$w^+/w^+$; P{20XUAS-IVS-GCaMP6f}attP40/P{TRiP.HMJ21280}attP40; P{R42F06-GAL4}attp2/P{TRiP.JF02847}attP2* |
| | *Rdl* RNAi | *$w^+/w^+$; P{20XUAS-IVS-GCaMP6f}attP40/+; P{R42F06-GAL4}attp2/P{UAS-Rdl RNAi8-10}* |
| **3B, 3C** | *HisCl1* RNAi | *$w^+/w^+$; P{R39H12-GAL4}su(Hw)attP5/P{TRiP.HMJ21280}attP40; P{VT50384-lexA}attP2, P{13XLexAop2-IVS-GCaMP6m p10}VK00005/P{TRiP.JF02847}attP2* |
| | *ort* RNAi | *$w^+/w^+$; P{R39H12-GAL4}su(Hw)attP5/P{VSH330348}attP40; P{VT50384-lexA}attP2, P{13XLexAop2-IVS-GCaMP6m-p10}VK00005* |
| **3D** | T4/T5ad > *GCaMP8m* | *$w^+/w^-$; +/+; P{T4/T5ad-GAL4}attP2, P{20XUAS-IVS-jGCaMP8m}VK00005/TM6B* |
| **3E–3G, S6, S7** | Control | *$w^+/w^-$; +/+; P{T4/T5ad-GAL4}attP2, P{20XUAS-IVS-jGCaMP8m}VK00005/+* |
| | *HisCl1* RNAi | *$w^+/w^-$; P{TRiP.HMJ21280}attP40/+; P{T4/T5ad-GAL4}attP2, P{20XUAS-IVS-jGCaMP8m}VK00005/P{TRiP.JF02847}attP2* |
| | *Rdl* RNAi | *$w^+/w^-$; +/+; P{T4/T5ad-GAL4}attP2, P{20XUAS-IVS-jGCaMP8m}VK00005/P{UAS-Rdl RNAi8-10}* |
| | *ort* RNAi | *$w^+/w^-$; P{VSH330348}attP40/+; P{T4/T5ad-GAL4}attP2, P{20XUAS-IVS-jGCaMP8m}VK00005/+* |
| | *Rdl* + *ort* RNAi | *$w^+/w^-$; P{VSH330348}attP40/+; P{T4/T5ad-GAL4}attP2, P{20XUAS-IVS-jGCaMP8m}VK00005/P{UAS-Rdl RNAi8-10}* |
| **5** | T4/T5 | *$w^+/w^-$; P{UAS-Dcr-2.D}2/+; P{R39H12-GAL4}attP2/+* |
| | *ort* RNAi | *$w^+/w^+$; P{VSH330348}attP40/+; +* |
| | T4/T5 > *ort* RNAi | *$w^+/w^-$; P{UAS-Dcr-2.D}2/P{VSH330348}attP40; P{R39H12-GAL4}attP2/+* |
| | *Rdl* RNAi | *$w^+/w^+$; +/+; P{UAS-Rdl RNAi8-10}/+* |
| | T4/T5 > *Rdl* RNAi | *$w^+/w^-$; P{UAS-Dcr-2.D}2/+; P{R39H12-GAL4}attP2/P{UAS-Rdl RNAi8-10}* |
| | *Rdl* + *ort* RNAi | *$w^+/w^+$; P{VSH330348}attP40/+; P{UAS-Rdl RNAi8-10}/+* |
| | T4/T5 > *Rdl* + *ort* RNAi | *$w^+/w^-$; P{UAS-Dcr-2.D}2/P{VSH330348}attP40; P{R39H12-GAL4}attP2/P{UAS-Rdl RNAi8-10}* |
| **S8** | Control | *P{hs-FLPG5.PEST}attP3/$w^-$; P{VT043070-p65.AD}attP40, P{UAS-FRT-stop-FRT-CD4tdGFP}/+; P{R39H12-DBD.GAL4}/+* |
| | *ort* RNAi | *P{hs-FLPG5.PEST}attP3/$w^-$; P{VT043070-p65.AD}attP40, P{UAS-FRT-stop-FRT-CD4tdGFP}/P{VSH330348}attP40; P{R39H12-DBD.GAL4}attP2/+* |
